# Flexible statistical methods for estimating and testing effects in genomic studies with multiple conditions

**DOI:** 10.1101/096552

**Authors:** Sarah M. Urbut, Gao Wang, Peter Carbonetto, Matthew Stephens

## Abstract

We introduce new statistical methods for analyzing genomic datasets that measure many effects in many conditions (e.g., gene expression changes under many treatments). These new methods improve on existing methods by allowing for arbitrary correlations in effect sizes among conditions. This flexible approach increases power, improves effect estimates, and allows for more quantitative assessments of effect-size heterogeneity compared to simple “shared/condition-specific” assessments. We illustrate these features through an analysis of locally-acting variants associated with gene expression (“cis eQTLs”) in 44 human tissues. Our analysis identifies more eQTLs than existing approaches, consistent with improved power. We show that while genetic effects on expression are extensively shared among tissues, effect sizes can still vary greatly among tissues. Some shared eQTLs show stronger effects in subsets of biologically related tissues (e.g., brain-related tissues), or in only one tissue (e.g., testis). Our methods are widely applicable, computationally tractable for many conditions, and available online.

## Introduction

Genomic studies often involve estimating and comparing many effects across multiple conditions or outcomes. Examples include studying changes in expression of many genes under multiple treatments^1^, differences in histone methylation at many genomic locations in multiple cell lines^2^, effects of many genetic variants on risk of multiple diseases^3^, and effects of expression quantitative trait loci (eQTLs) in multiple cell-types or tissues^4–6^. Analyses often aim to identify “significant” non-zero effects, and to identify differences in effects among conditions (e.g., tissue-specific effects), which may yield biological insights.

A common analysis strategy for such studies is to separately analyze each condition in turn, then compare the “significant” results among conditions. Although appealingly simple, this “condition-by-condition” approach is unsatisfactory in several ways: it under-represents sharing of effects among conditions because shared effects will be insignificant in some conditions by chance, and it misses the power gains that come from sharing information across conditions^5^.

To address deficiencies of condition-by-condition analyses, several groups have developed methods for *joint* analysis of multiple conditions^2–3,5–15^. The simplest methods build on traditional meta-analysis methodology^8,9^, and assume that non-zero effects are shared across all conditions. Other methods are more flexible, allowing for condition-specific effects, for sharing of effects among subsets of conditions, and for heterogeneity in the shared effects^5,6,12^. The most flexible methods adapt themselves to each dataset by learning patterns of sharing using a hierarchical model^5^.

Nonetheless, existing methods have important limitations. First, they make restrictive assumptions about the correlations of non-zero effects among conditions. For example, Flutre *et al*.^5^ assume these correlations are non-negative and equal. Correlations may be negative in some applications; e.g., genetic variants that increase one trait may decrease another. And some conditions may be more correlated than others; for example, in our eQTL application (below), brain tissues are strongly correlated with one another. Second, the most flexible methods are computationally intractable for moderate numbers of conditions (e.g., 44 tissues in our eQTL application). Existing solutions to this problem substantially reduce flexibility. For example, Flutre *et al*.^5^ solve the computational problem by restricting effects to be shared in all conditions, or to be specific to a single condition. Alternatively, Wei *et al*.^12^ allow for all possible patterns of sharing, but only under the restrictive assumption that non-zero effects are uncorrelated among conditions. Third, existing methods typically focus only on *testing* for significant effects in each condition, and not on *estimating effect sizes* which, as we illustrate, is important for assessing heterogeneity among conditions.

We introduce more flexible statistical methods that combine the most attractive features of existing approaches, while overcoming their major limitations. The methods, which we refer to as “multivariate adaptive shrinkage” (`mash`), build on recent approaches^16^ for testing and estimating effects in a *single* condition, extending these approaches to *multiple* conditions. Key features of `mash` include: (i) it is *flexible*, allowing for both shared and condition-specific effects, and arbitrary patterns of correlation among conditions; (ii) it is *computationally tractable* for hundreds of thousands of tests in dozens of conditions, or more; (iii) it provides not only measures of significance, but also *estimates of effect sizes*, together with measures of uncertainty; (iv) it is *adaptive*, meaning that its behavior adapts to the patterns present in the data; and (v) it is *generic*, requiring only a matrix containing the observed effects in each condition and a matrix of the corresponding standard errors. (Alternatively, `mash` can be supplied with just matrix of *Z* scores, although this reduces the ability to estimate effect sizes.) Together, these features make `mash` the most flexible and widely applicable method available for estimating and testing multiple effects in multiple conditions.

To demonstrate the potential for `mash` to provide novel insights, we apply it to analyze *cis* eQTL effects in 16,069 genes across 44 human tissues. Focusing on the strongest *cis* eQTLs, we find that while most eQTLs are shared by many tissues, effect sizes can still vary considerably. Our results suggest that when assessing effects that are “tissue-specific” versus “tissue-consistent”, we should pay careful attention to the *sizes* of effects, and not only the tests for significance.

## Results

### Methods overview

Our method, `mash`, estimates effects of many “units” (*J*) in many conditions (*R*), allowing that effects may be sparse (*i.e*., many zero effects), and allowing for correlations among non-zero effects in different conditions. For example, in multi-tissue eQTL studies, the “units” are eQTLs (*J* > 10,000), the conditions are different tissues (*R* = 44), and `mash` estimates the effect of each eQTL in each tissue, allowing for cross-tissue sharing and tissue-specificity of eQTLs.

To apply `mash`, the analyst must first conduct a condition-by-condition analysis to obtain an effect estimate and corresponding standard error for each unit in each condition. These estimates are the inputs for a two-step Empirical Bayes (EB) procedure: (1) learn patterns of sparsity, sharing and correlations among effects from the condition-by-condition results; (2) combine these learned patterns with the condition-by-condition results to produce improved effect estimates and corresponding measures of significance. These steps are summarized here, and in Fig. 1. (See Methods for more details.)

**Figure 1.**
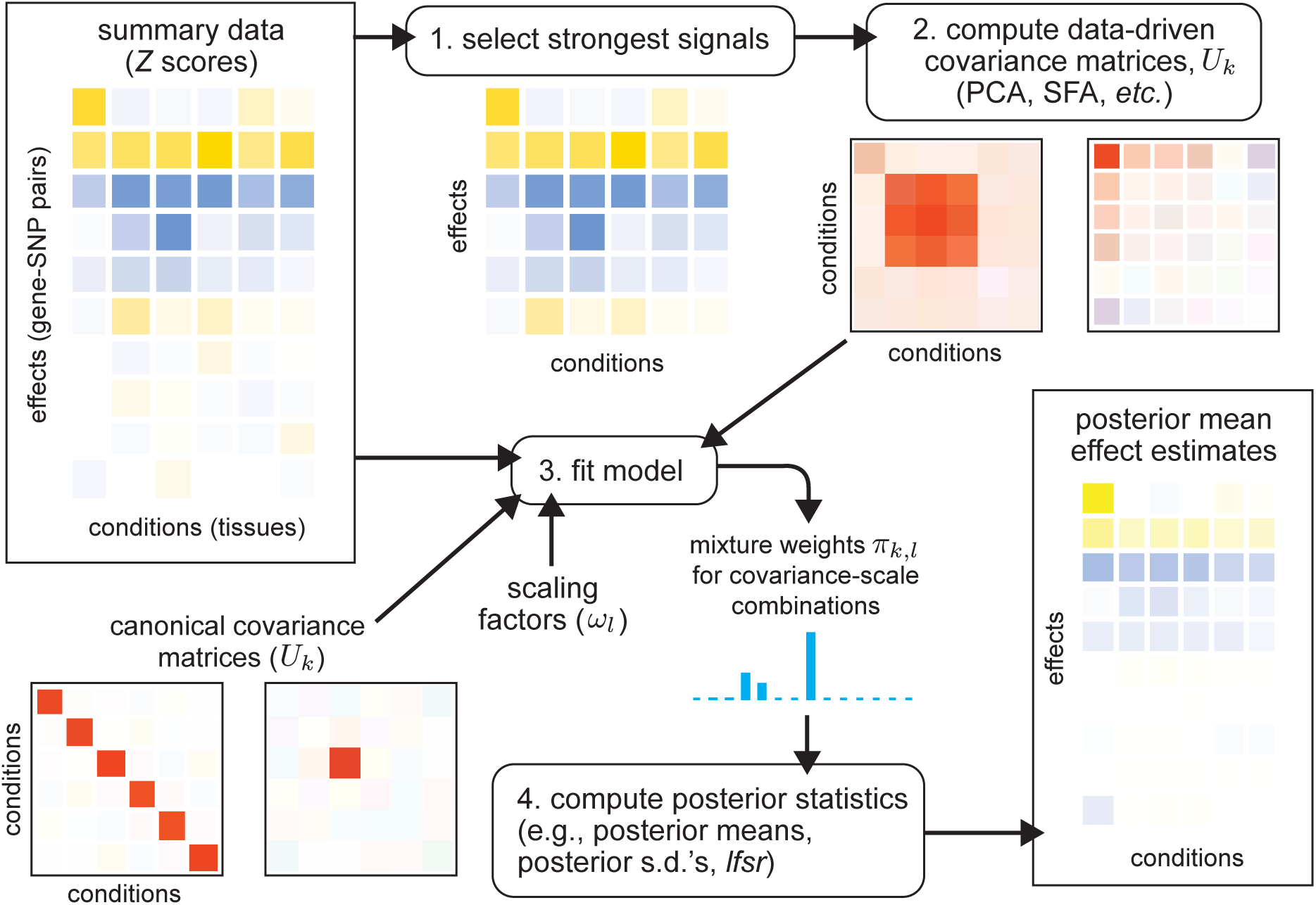
Overview of fitting procedure in mash, which estimates the multivariate distribution of effects present in the data. The data consist of a matrix of summary data (e.g., *Z* scores) for a large number of units (e.g., gene-SNP pairs) in multiple conditions (e.g., tissues), and, optionally, their standard errors (not shown). Color indicates the sign (positive, negative) of an effect (blue, yellow) or covariance (blue, red), with shading intensity indicating size. After selecting rows containing the strongest signals (**1**)—in this example, the top 6 rows—we apply covariance estimation techniques to estimate candidate “data-driven” covariance matrices *U*_*k*_ (**2**). To these, we add “canonical” covariance matrices *U*_*k*_, including the identity matrix, and matrices representing condition-specific effects. Each covariance matrix represents a pattern of effects that may occur in the data. We scale each covariance matrix by a grid of scaling factors, *ω*_*l*_, varying from “very small” to “very large”, which allows for *a priori* effect sizes to range from very small to very large. Using the entire data set, we compute maximum-likelihood estimates of the weights (relative frequencies) *π*_*k*, *l*_ for each (*U*_*k*_, *ω*_*l*_) combination (**3**), thereby learning how commonly each pattern-effect size combination occurs in the data. Finally, we compute posterior statistics using the fitted model (**4**); the posterior mean estimates shown in the bottom-right illustrate that effect estimates are “shrunk” adaptively using the fitted `mash` model.

In brief, let *b* denote the vector of true effects for a single unit across *R* conditions. We capture correlations and sharing of effects among conditions using a mixture model,

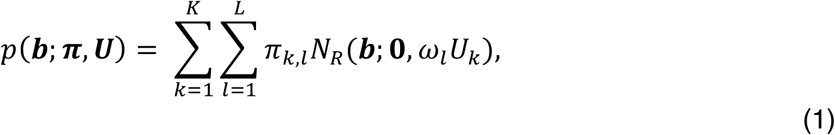

where *N*_*R*_(·; *μ*, Σ) denotes the multivariate normal density in *R* dimensions with mean *μ* and variance-covariance matrix Σ; each *U*_*k*_ is a covariance matrix that captures a pattern of effects; each *ω*_*l*_. is a scaling coefficient that corresponds to a different effect size; and the mixture proportions *π*_*k*, *l*_ determine the relative frequency of each covariance-scale combination. The scaling coefficients *ω*_*l*_. take values on a fixed dense grid that spans from “very small” to “very large”, capturing the full range of possible effects.

Step 1 of `mash` learns patterns of sparsity, sharing and correlations by estimating covariance matrices *U* and mixture proportions *π* in two sub-steps:

*Step 1a:* Generate candidate covariance matrices *U* = (*U*_1_,…, *U*_*K*_). This list includes both “data-driven” matrices that are estimated from the strongest signals in the condition-by-condition results (see Methods), and “canonical” matrices that have simple interpretations. For example, the “canonical” matrices include the identity matrix (representing independent effects across all conditions); a matrix of all ones (representing effects that are equal in all conditions); and *R* matrices that represent effects that are specific to each condition.
*Step 1b:* Given *U*, estimate *π* by maximum likelihood from the condition-by-condition results. This step can rescue imperfections in Step 1a by assigning low weight to covariance matrices that are not well-supported by the data. This step also adapts to sparse effects; if most effects are zero, or small, this step will put most weight on small effects (*i.e*., small scaling coefficients, *ω*).

Step 2 of `mash` uses Bayes’ theorem to compute the posterior distribution for each effect given the condition-by-condition results and the fitted prior (eq. 1). These posterior distributions yield improved effect estimates—posterior means and standard deviations—that account for sparsity and correlations among effects. We use these estimates to define quantitative measures of effect sharing between any two conditions: “sharing by sign” (effects have the same sign), and “sharing by magnitude” (effects have similar magnitude—here defined to be within a factor of 2, although other thresholds could be used; see Supplementary Note). The posterior distributions also yield a condition-specific measure of significance for each effect, the “local false sign rate”, or *lfsr*^16^, that is analogous to a false discovery rate, but more stringent because it requires true discoveries to be not only non-zero, but also correctly signed. Finally, `mash` also computes, for each unit, a Bayes Factor that summarizes the overall significance of the unit—*i.e*., overall evidence for a non-zero effect in *any* condition.

### Simulation studies

We ran simulations to compare `mash` with existing methods. We simulated effects for 20,000 units in 44 conditions, with 400 units having non-null effects. Non-null effects were simulated under two different scenarios:

1. “Shared, structured effects”: Effects are largely shared (non-null units have an effect in many conditions), and “structured”—that is, similar in size and direction, with greater similarity among some subsets of conditions. This scenario was based on the fit of the model (eq. 1) to the eQTL data (see Methods).
2. “Shared, unstructured effects”: Effects are shared (non-null units have an effect in every condition) but unstructured (independent) across conditions.

We used these simulations to compare `mash` with several other methods: `mash-bmalite`, a simplified version of `mash` that uses only the BMAlite models from Flutre *et al*^5^; `ash`^16^, a univariate analogue of `mash` that analyzes each condition separately; and `metasoft`^9,14^, which implements several multivariate tests corresponding to different models (the “fixed effects”, or FE^9^, model, which assumes equal effects in all conditions, the RE2 model^9^, which allows for normally-distributed variation in effects among conditions, and the BE model, which extends the RE2 model to allow for effects are exactly zero^14^).

The simulation results are summarized in Fig. 2 and Supplementary Table 1. We assessed the methods on three important tasks: distinguishing non-null units from null units (Fig. 2a–b); distinguishing non-null effects (*i.e*., non-null unit-condition pairs) from null effects (Fig. 2c–d); and estimating effect sizes (Fig. 2e–f). (We included metasoft results for the first task only because it does not provide multivariate effect estimates, and because it is unclear how best to combine unit-level significance (*p* value) and condition-specific significance (*m* value) into a significance measure for each unit-condition pair.)

**Figure 2.**
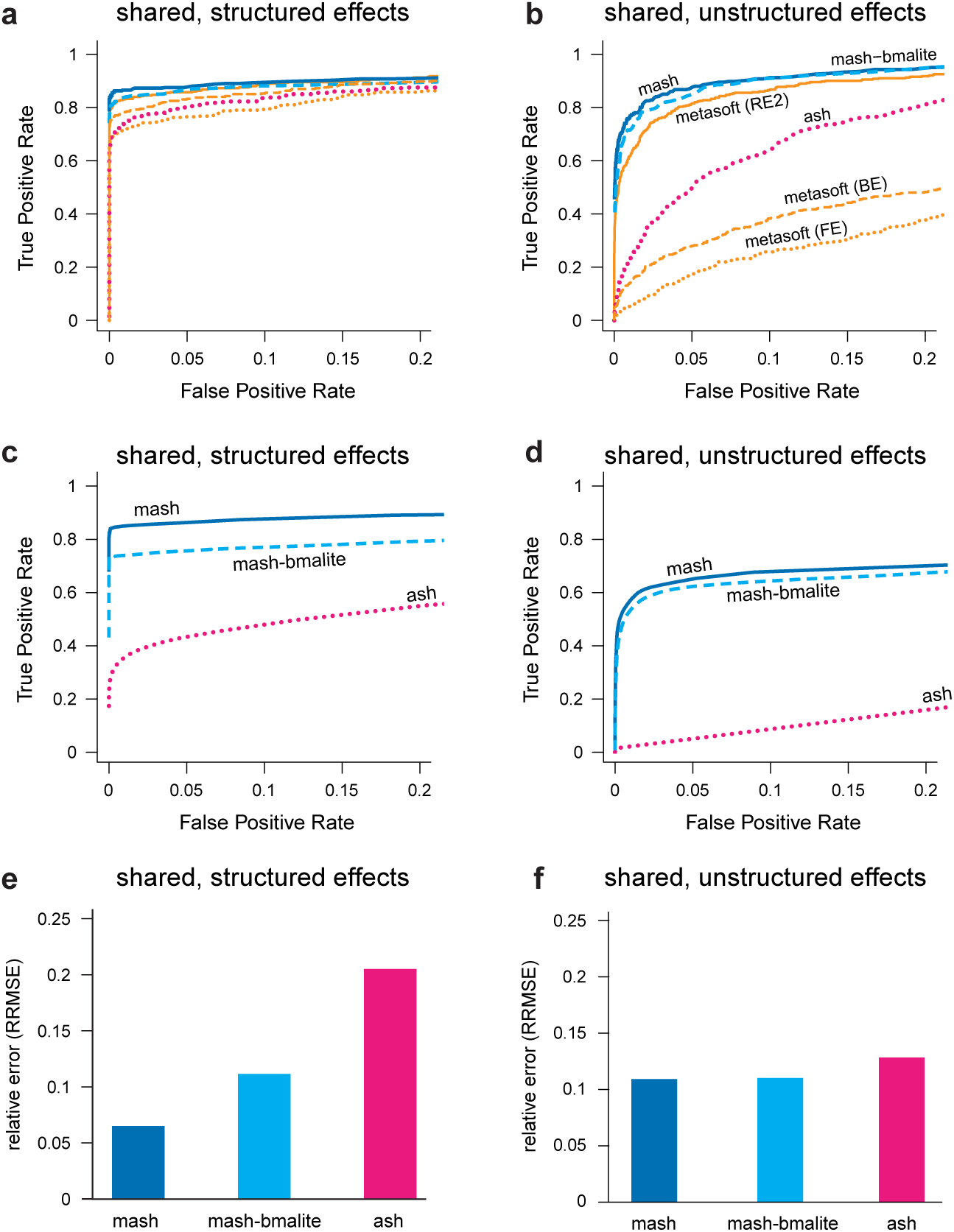
Comparison of methods on simulated data. Results are shown for two simulation scenarios: “shared, structured effects”, in which the non-zero effects are shared among conditions in complex, structured ways similar to patterns of eQTL sharing in the GTEx data; and “shared, unstructured effects”, in which the non-zero effects are shared among conditions but independent. Each simulation result involves *n* = 20,000 independent units observed at *R* = 44 conditions, with 400 non-null units. Panels **a–b** show ROC curves for detecting significant units (*n* = 20,000 discoveries), based on unit-specific measures of significance (as in traditional meta-analyses). Panels **c-d** show ROC curves for detecting significant effects (*n* × *R* = 44 × 20,000 = 880,000 discoveries), which requires effect-specific measures of significance. In **c–d**, we also require the estimated sign (+/–) of each significant effect to be correct to be considered a “true positive”. Panels **e** and **f** summarize the error in the estimated effects relative to the error from a simple condition-by-condition analysis (Relative Root Mean Squared Error, or RRMSE for short). Our new method (`mash`) outperformed other methods, particularly in the “shared, structured effects” scenario.

In the “shared, structured” scenario, `mash` markedly outperformed the other methods in all three tasks (Fig. 2, Supplementary Table 1). This is expected because `mash` is the only method that attempts to fully exploit the structure in these effects. Further, `mash` found essentially all (>99%) of the signals that the other methods found, plus additional signals (Supplementary Table 2).

In the “shared, unstructured” scenario, `mash` and `mash`-bmalite are the best methods, finding almost exactly the same signals (Supplementary Table 2), with `metasoft`-RE2 close behind as it was applied to the task it was designed for (testing unit-level significance). The strong similarity in performance between `mash` and `mash-bmalite` is explained by `mash` putting negligible weight on the data-driven covariance matrices, which are unnecessary in this scenario. This illustrates `mash`’s ability to adapt to the data, increasing power in complex scenarios without being unduly penalized in simpler scenarios.

Note that `mash` uses two distinct strategies to improve accuracy of effect estimates: (i) it shrinks estimates towards zero, which improves average accuracy because most effects are null; (ii) in the presence of “structured effects”, it shares information across conditions to improve accuracy. For example, if a unit has effects that are similar across a subset of conditions, averaging the effect estimates in those conditions will improve accuracy. Both these strategies contribute to the strong performance of `mash` in the “shared, structured” scenario.

These simulations are meant to be illustrative rather than comprehensive. Indeed, although numerical comparisons can help choose between methods, we believe that *qualitative* factors are equally important. For example, the fact that `mash` produces effect size estimates is a key feature that distinguishes it from most existing methods. See Supplementary Note for a detailed discussion on connections and differences between the methods.

### GTEx *cis*-eQTL analysis

To illustrate `mash` in a substantive application, we analyzed eQTLs across 44 human tissues, using data from the Genotype Tissue Expression (GTEx) project^17^. The GTEx project aims to provide insights into the mechanisms of gene regulation by studying gene expression and regulation in multiple tissues from human donors. One fundamental question is which SNPs are eQTLs (*i.e*., associated with gene expression) in which tissues. Answering this could help distinguish regulatory mechanisms that are tissue-specific or shared, and, by integrating with results from genome-wide association studies (GWAS), could help identify the most relevant tissues for complex disease^17,18^.

As input, `mash` requires results from a condition-by-condition analysis (effect estimates and standard errors). We used the effect estimates and standard errors for candidate local (“*cis*”) eQTLs for each gene (GTEx data release 6). These were obtained by performing (univariate) single-SNP eQTL analyses in each tissue using Matrix eQTL^19^. Expression levels were corrected for population structure (using genotype principal components^20^) and other confounding factors (both measured factors such as age and sex, and unmeasured factors estimated using factor analysis^21^), then rank-transformed to the corresponding quantiles of a standard normal distribution. Because these estimates were obtained by single-SNP analysis, the estimated effects for each SNP reflect both the effects of the SNP itself *and SNPs in LD with it*. Thus, our analyses do not distinguish causal eQTLs from those that are in LD with the causal eQTLs; see Discussion and Supplementary Note.

We analyzed the 16,069 genes for which effect estimates were available for all 44 tissues considered; the filtering criteria used by GTEx ensure that these genes show some expression in all 44 tissues.

### mash improves model fit

To assess the improved fit of `mash` compared with the simpler `mash-bmalite`, we used cross-validation; we fitted each model to a random subset of units (“training set”), and assessed fit by the log-likelihood on the remaining units (“test set”). We found that `mash` improved the test set log-likelihood very substantially (by 23,796; Supplementary Fig. 1). Further, `mash` placed 79% of the weight on the data-driven covariance matrices. These results confirm that our methods for estimating data-driven covariance matrices are sufficiently effective that they better capture most effects than do the canonical matrices used by existing methods.

### Identification of data-driven patterns of sharing

The data-driven covariance matrix with largest `mash` weight is shown in Fig. 3 (34% weight). This covariance matrix captures several important features: (i) effects are positively correlated among all tissues; (ii) the brain tissues—and, to a lesser extent, testis and pituitary—are particularly strongly correlated with one another, and less correlated with other tissues; (iii) effects in whole blood tend to be less correlated with other tissues. Many other data-driven covariance matrices estimated by `mash` also have positive correlations among all tissues and/or highlight heterogeneity between brain tissues and other tissues, confirming these as common features of these data (Supplementary Fig. 2). Other components capture less prevalent patterns, such as effects that are appreciably stronger in one tissue (e.g., Supplementary Fig. 2b).

**Figure 3.**
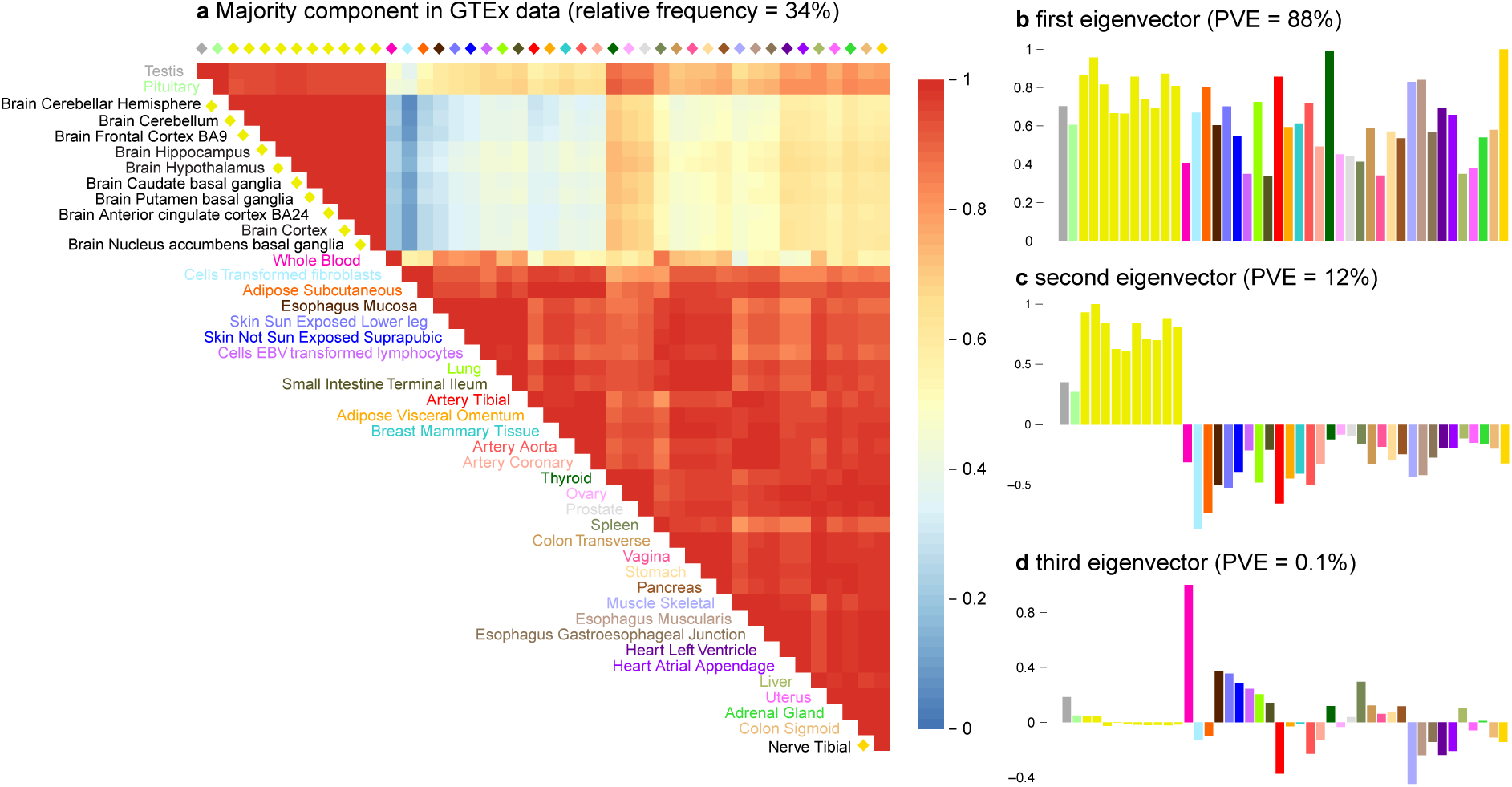
Summary of primary patterns identified by mash in GTEx data. Shown are the heatmap of the correlation matrix (**a**) and bar plots of the first three eigenvectors (**b, c, d**) of the covariance matrix *U*_*k*_ corresponding to the dominant mixture component identified by `mash` (*n* = 16,069 independent gene-SNP pairs). This component accounts for 34% of all weight in the GTEx data. Tissues are color-coded as indicated by the tissue labels in the heatmap. The first eigenvector (**b**) reflects broad sharing among all tissues, with all effects in the same direction; the second eigenvector (**c**) captures differences between brain (and, to a lesser extent, testis and pituitary) and other tissues; the third eigenvector (**d**) primarily captures effects that are stronger in whole blood.

### Patterns of sharing inform effect size estimates

Having estimated patterns of sharing, `mash` exploits these patterns to improve effect estimates at each candidate eQTL. Although we cannot directly demonstrate improved accuracy of effect estimates in the GTEx data (for this, see simulations above), individual examples can provide insight into how `mash` achieves improved accuracy. Figure 4 shows three illustrative examples, which we discuss in turn.

**Figure 4.**
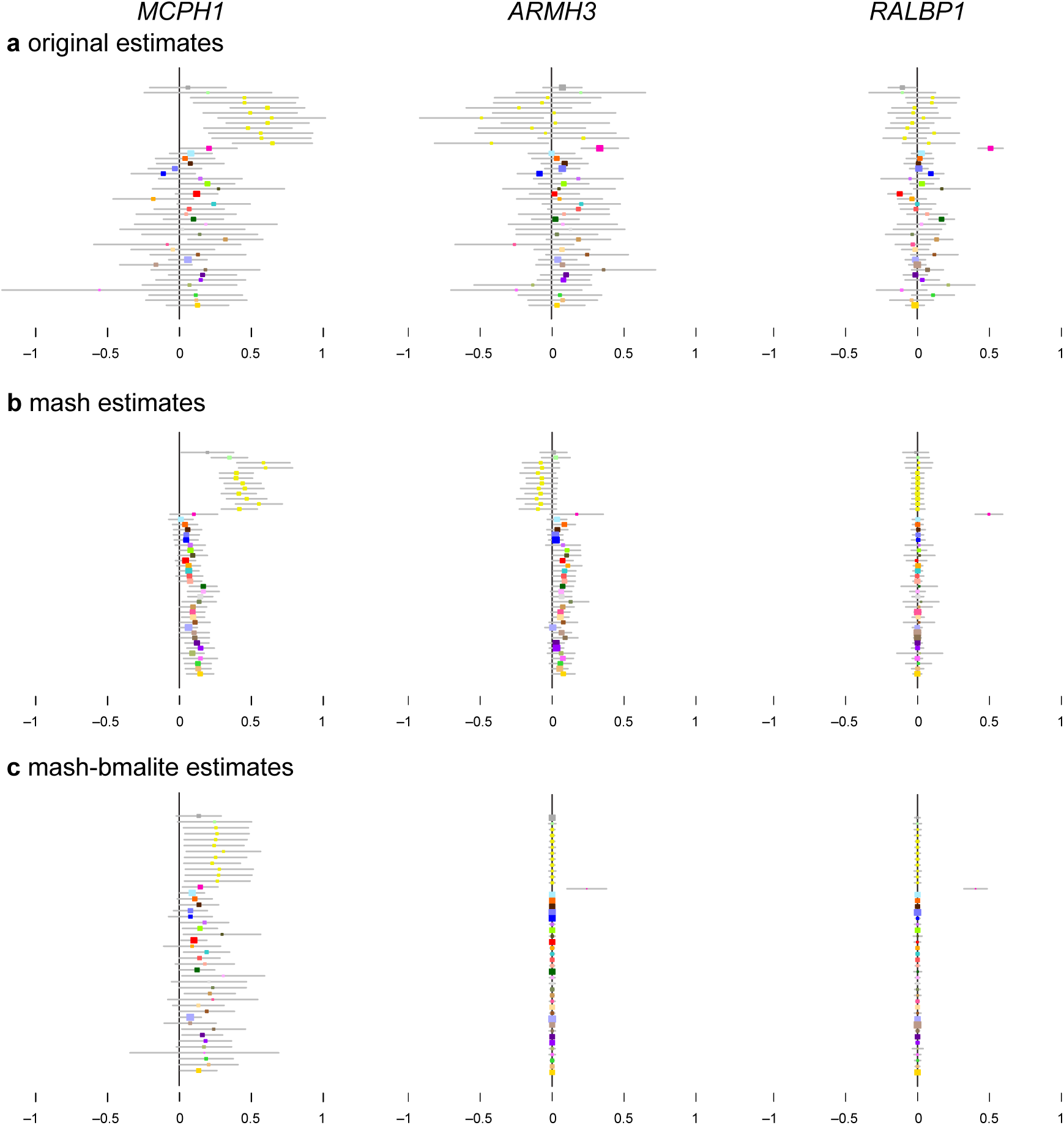
Examples illustrating that mash uses learned patterns of sharing to inform effect estimates in the GTEx data. In panel **a**, each colored dot shows the original (“raw”) effect estimate for a single tissue (color-coded as in Fig. 3), with grey bars indicating ±2 standard errors. These are the data provided to `mash`. Panel **b** shows the corresponding `mash` estimates. In each case, `mash` combines information across all tissues, using the background information (patterns of sharing) learned from data on all eQTLs to produce more precise estimates. Panel **c** shows, for contrast, the corresponding estimates from `mash`-bmalite, which, due to its more restricted model, fails to capture features clearly apparent in the original data, such as strong brain effects in *MCPH1*. In **b** and **c**, colored dots are posterior means, and error bars depict ±2 posterior standard deviations. For all estimates, *n* = 83–430 individuals, depending on the tissue (Supplementary Table 3).

In the first example (Fig. 4, left-hand column), raw effect estimates in each tissue are mostly positive, and strongest in brain tissues. The `mash` estimates are *all* positive: the few modest negative estimates are outweighed by the strong background information that effects are highly correlated among tissues and mostly positive. This weighing of evidence was done using Bayes’ rule. Humans are notoriously bad at weighing background information against specific instances—they tend to underweight background information when presented with specific data^22^—so this behavior may or may not be intuitive. The `mash` effect estimates are also appreciably larger in brain tissues than in other tissues. This is a result of using Bayes’ rule to combine the effect estimates at *this* eQTL with the background information on heterogeneity of brain and non-brain effects learned from *all* eQTLs.

In the second example (Fig. 4, middle column), most effect estimates in non-brain tissues are positive (30 out of 34), but modest in size, and only one is nominally significant (*p* < 0.05). However, by combining information among tissues, `mash` estimated that all effects in non-brain tissues are positive, and mostly “significant” (*lfsr* < 0.05). By contrast, the estimated effects in brain tissues are inconsistent (both positive and negative) and so `mash` is not confident about the sign of effects in brain tissues. This example illustrates that `mash` can learn to treat subsets of conditions differently; `mash` learned that effects in brain tissues are occasionally different from effects in other tissues, and therefore did not draw strong inferences in the brain based on the other tissues.

In the third example (Fig. 4, right-hand column), the effect estimates vary in sign, and are modest except for a very strong signal in whole blood. While whole-blood-specific effects were estimated to be rare, `mash`—again, using Bayes’ theorem—recognized that the strong signal at this eQTL outweighs the background information, and estimated a strong effect in blood with insignificant effects in other tissues. This illustrates that `mash`, although focused on combining information across tissues, can still recognize—and shed light on—tissue-specific patterns when they exist.

In all three examples, the `mash` estimates are noticeably different from those obtained by `mash-bmalite` (Fig. 4c) and ash (Supplementary Fig. 3). The decreased flexibility of `mash-bmalite` compared to `mash` is particularly evident in the first example where `mash-bmalite` estimates are almost constant across tissues, hiding the stronger effects in brain-related tissues that are clearly visible in the original data, and maintained by `mash`.

### Increased identification of significant effects

Consistent with our simulation results, `mash` identified many more significant effects than either `mash-bmalite` or ash. To avoid double-counting eQTLs in the same gene that are in LD with one another, we assessed the significance of only the “top” SNP in each gene (the SNP with the largest univariate |*Z*|-statistic across tissues). Thus, our results are based on 16,069 candidate eQTLs, each with effect estimates in 44 tissues, for a total of 44 × 16,069 = 707,036 effects.

The vast majority of top SNPs show a strong signal in at least one tissue (97% have a maximum |*Z*| score exceeding 4), consistent with most genes containing at least one eQTL in at least one tissue. However, in the tissue-by-tissue analysis ash identified only 13% of these effects as “significant” at *lfsr* < 0.05; that is, the univariate analysis was highly confident in the direction of the effect in only 13% of cases. By comparison, `mash-bmalite` identified 39% as significant at the same threshold, and `mash` identified 47%. Similar to our simulations, the significant associations identified by `mash` include the vast majority (95%) of those found by either of the other methods (Supplementary Table 4).

Overall, `mash` found 76% (12,171 out of 16,069) of the top SNPs to be significant in at least one tissue. We refer to these as the “top eQTLs” in subsequent sections.

### Sharing of effects among tissues

To investigate sharing of the top eQTLs among tissues, we assessed sharing of effects by sign and by magnitude (effects have the same sign and are within a factor of 2 in size). Because of the large differences between brain and non-brain tissues, we also show results separately for these subsets. The results (Table 1, Fig. 5) confirm extensive eQTL sharing among tissues, particularly among brain tissues. Sharing by sign always exceeds 85%, and is as high as 96% among brain tissues. (Furthermore, these numbers may underestimate the sharing by sign of actual causal effects due to the impact of multiple eQTLs in LD; see Supplementary Note and Supplementary Fig. 4.) Sharing by magnitude is necessarily lower because sharing by magnitude implies sharing by sign. On average, 36% of tissues show an effect within a factor of 2 of the strongest effect at each top eQTL. However, within brain tissues this increases to 76%. Thus, not only do eQTLs tend to be shared among brain tissues, but effect sizes tend to be homogeneous. Because our analyses were based on the top eQTLs, our results reflect patterns of sharing only among stronger *cis* eQTLs; weaker eQTLs may show different patterns.

**Figure 5.**
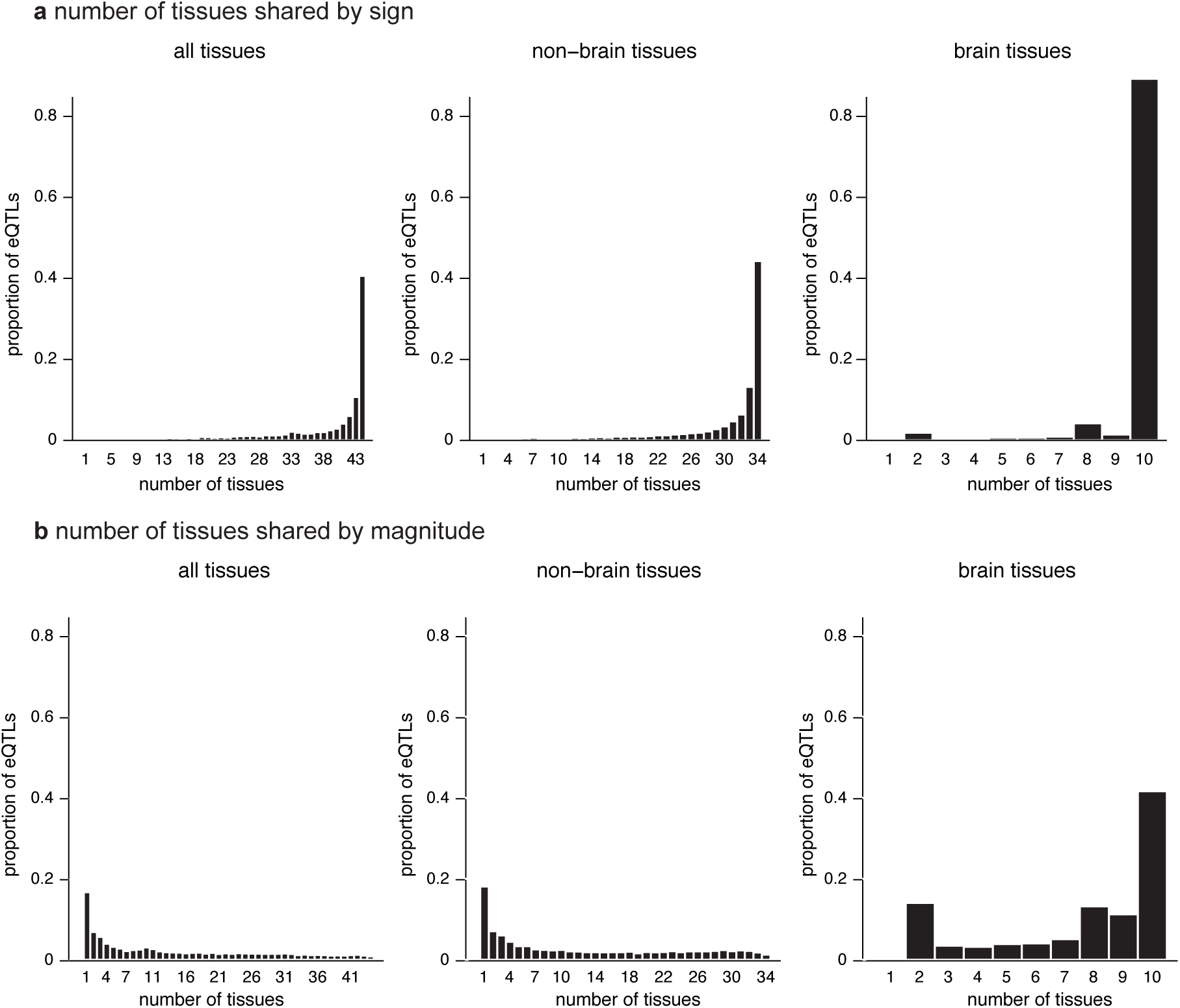
Number of tissues shared by sign and magnitude. Histograms show estimated number of tissues in which top eQTLs are “shared,” considering all tissues (*n* = 12,171 gene-SNP pairs with a significant eQTL in at least one tissue), non-brain tissues (*n* = 12,117), and brain tissues only (*n* = 8,474), and using two different sharing definitions, by sign (**a**) and by magnitude (**b**). Sharing by sign means that the eQTLs have the same sign in the estimated effect; sharing by magnitude means that they also have similar effect sizes (within a factor of 2).

**Table 1.** Summary of sharing among top eQTLs. Numbers give the proportion of effects meeting a given sharing criterion: “shared by sign” requires that the effect have the same sign as the strongest effect across tissues; “shared by magnitude” requires that the effect also be within a factor of 2 of the strongest effect. Numbers in parentheses were obtained from secondary `mash` analyses conducted separately in brain and non-train tissues.

| sharing criterion | proportion of effects |  |  |
| --- | --- | --- | --- |
|  | all tissues | non-brain | brain |
| shared by sign | 0.85 | 0.85 (0.88) | 0.96 (0.98) |
| shared by magnitude | 0.36 | 0.40 (0.44) | 0.76 (0.85) |

Some tissues share eQTLs more than others. Fig. 6 summarizes eQTL sharing by magnitude between all pairs of tissues (see Supplementary Fig. 5 for sharing by sign). In addition to strong sharing among brain tissues, `mash` identified increased sharing among other biologically related groups, including arteries (tibial, coronary and aortal), two groups of gut tissues (one group containing esophagus and sigmoid colon, the other containing stomach, terminal ilium of the small intestine and transverse colon), skin (sun-exposed and non-exposed), adipose (subcutaneous and visceral-omentum) and heart (left ventricle and atrial appendage). Figure 6 also reveals heterogeneity in effect sizes in cerebellum versus non-cerebellum tissues, and highlights sharing between pituitary and brain tissues.

**Figure 6.**
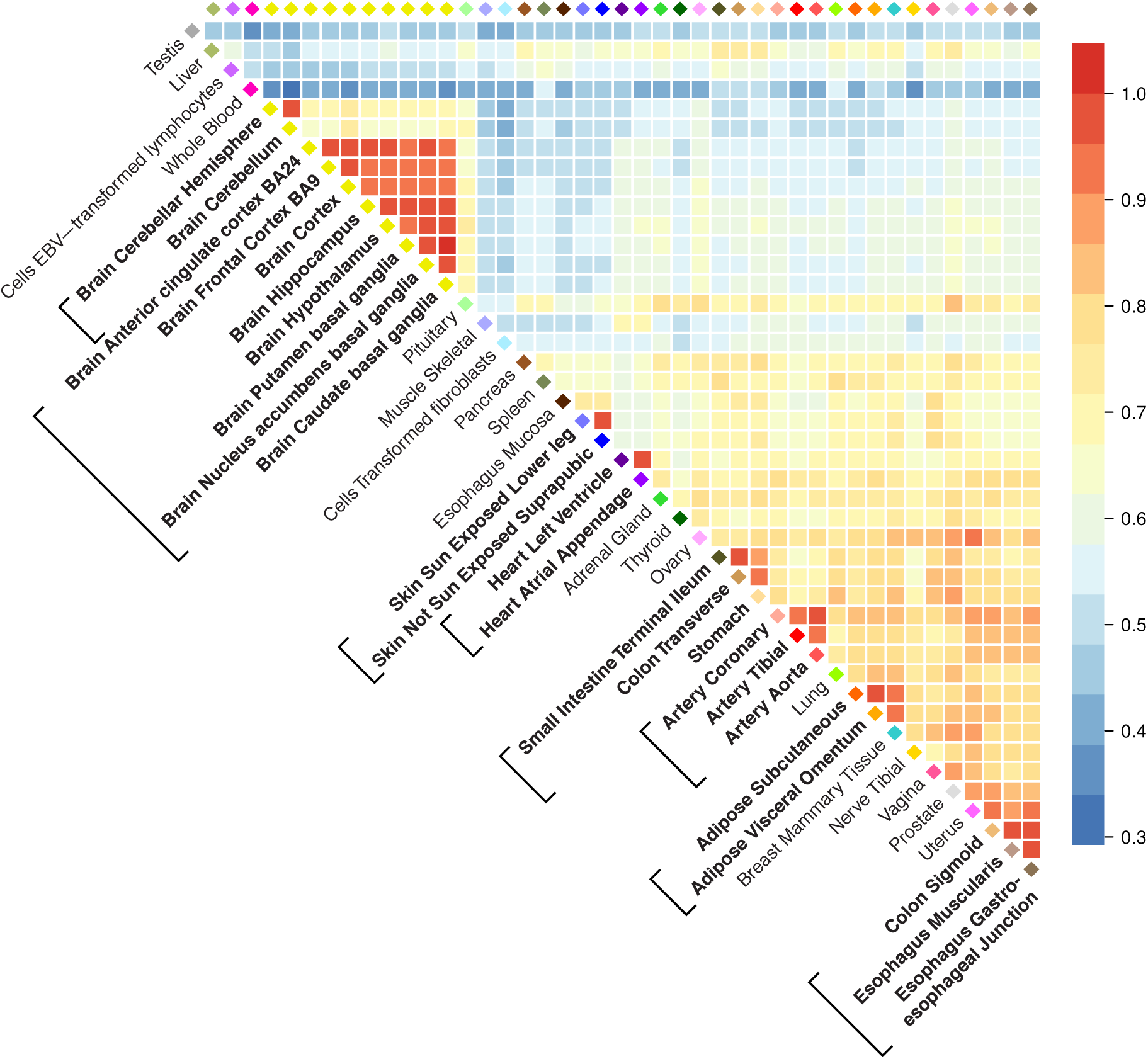
Pairwise sharing by magnitude of eQTLs among tissues. For each pair of tissues, we considered the top eQTLs that were significant (*lfsr* < 0.05) in at least one of the two tissues, and plotted the proportion of these that are “shared in magnitude”—that is, have effect estimates that are the same sign and within a factor of 2 in size of one another (*n* = 5,605–9,811 gene-SNP pairs, depending on pair of tissues compared). Brackets surrounding tissue labels highlight groups of biologically related tissues mentioned in the text as showing particularly high levels of sharing.

Different levels of effect sharing among tissues means that effect estimates in some tissues gain more precision than others from the joint analysis. To quantify this, we computed an “effective sample size” (ESS) for each tissue that reflects typical precision of its effect estimates (Supplementary Fig. 6, Supplementary Note). The ESS values are smallest for tissues that show more “tissue-specific” behaviour (e.g., testis, whole blood), and are largest for coronary artery, reflecting its stronger correlation with other tissues.

### Tissue-specific eQTLs

Despite high average levels of sharing of eQTLs among tissues, `mash` identified some eQTLs that are relatively “tissue-specific”. Indeed, the distribution of the number of tissues in which an eQTL is shared by magnitude has a mode at 1 (Fig. 5), representing a subset of eQTLs that have a much stronger effect in one tissue than in any other (henceforth, “tissue-specific” for brevity). Breaking down this group by tissue identified testis as the tissue with the most tissue-specific effects (Supplementary Fig. 7). Testis and whole blood stand out as having lower pairwise sharing of eQTLs with other tissues (Fig. 6). Other tissues showing larger than average tissue specificity (Fig. 5, Supplementary Fig. 7) include skeletal muscle, thyroid and transformed cell lines (fibroblasts and LCLs).

One possible explanation for tissue-specific eQTLs is tissue-specific expression; that is, if a gene is strongly expressed only in one tissue, this could cause an eQTL for that gene to show a strong effect only in that tissue. Whether or not a tissue-specific eQTL is due to tissue-specific expression could considerably impact biological interpretation. To assess whether tissue-specific eQTLs could be explained by tissue-specific expression we took genes with tissue-specific eQTLs and examined the distribution of expression in the eQTL-affected tissue relative to expression in other tissues. We found this distribution to be similar to genes without tissue-specific eQTLs (Supplementary Fig. 8). Therefore, most tissue-specific eQTLs identified here do not solely reflect tissue-specific expression.

## Discussion

The statistical benefits of joint multivariate analyses compared with univariate analyses are well documented, and increasingly widely appreciated. But we believe this potential nonetheless remains under-exploited. With `mash` we aim to provide a set of flexible and general tools to facilitate such analyses. In particular, `mash` is *generic* and *adaptive*. It is generic in that it can take as input any matrix of *Z* scores (or, preferably, a matrix of effect estimates and a matrix of corresponding standard errors) that test many effects in many conditions. These scores could come from many sources; e.g., from linear regression, generalized linear models or linear mixed models^13^. And `mash` is adaptive in that it learns correlations among effects from the data, allowing it to maximize power and precision for each setting. Consequently, `mash` should be widely applicable to many settings involving estimation of multivariate effects.

The `mash` method uses EB hierarchical modelling, and so is related to other EB methods^5,6,12^. Indeed, the `mash` framework essentially includes these methods as special cases (as well as simpler methods such as “fixed effects” and “random effects” meta-analyses^9,23^). One key feature that distinguishes `mash` from previous methods is that `mash` puts greater emphasis on *quantitative* estimation and assessment of effects. Moving away from binary-based models has at least two advantages. First, allowing for all possible binary configurations can create computational challenges. Second, in practice we have found that effects are often shared broadly among many conditions, and in such cases binary-based methods tend to infer that effects are non-zero in most or all conditions, even when the signal is modest in some conditions. This conclusion may be technically correct—for example, in our GTEx analysis it is not impossible that all eQTLs are somewhat active in all tissues. However, as our analysis has illustrated, a more quantitative focus can reveal variation in effect sizes that may be of considerable biological importance.

One limitation of our eQTL analysis is that, like most eQTL analyses, it does not distinguish between causal associations and those that are due to LD. This issue is particularly important to appreciate when, for example, cross-referencing GWAS associations with eQTL effect estimates; a GWAS-associated SNP may be a “significant” eQTL simply because it is in LD with another causal SNP. For single-tissue eQTL mapping, this problem has been addressed in several ways, including fine-mapping^24–29^ and co-localization^30–32^. For multi-tissue analysis, only more limited attempts exist to address this problem. For example, eQTLBMA^5^ implements a Bayesian approach to fine-mapping under the simplifying assumption that there is at most one causal SNP per gene^24,25^. However, this assumption becomes less plausible in analyses of many tissues, and developing more flexible multi-tissue fine-mapping methods seems an important future direction.

Dealing with multiple tests is often described as a “burden”. This likely originates from the fact that controlling family-wise error rate requires more stringent thresholds as the number of tests increases. However, modern analyses prefer to control the FDR^33^, which does not depend on the number of tests^34^. Consequently, the term “burden” is inaccurate and unhelpful. We believe that the results of many tests in many conditions should instead be viewed as an *opportunity*—an opportunity to learn about relationships among underlying effects, and make data-driven decisions to improve inferences. This will inevitably, it seems, involve modelling assumptions, and the challenge is to design flexible models that work well in a range of settings. The methods presented here represent a major step towards this goal.

URLs. Multivariate adaptive shrinkage (“`mash`”) software, https://github.com/stephenslab/mashr; code and data resources for GTEx analysis, https://github.com/stephenslab/gtexresults, https://doi.org/10.5281/zenodo.1296399; GTEx project, http://gtexportal.org; adaptive shrinkage (“ash”) software, https://github.com/stephens999/ashr; Sparse Factor Analysis (SFA) software, http://stephenslab.uchicago.edu/software.html; Extreme Deconvolution software, https://github.com/jobovy/extreme-deconvolution; METASOFT software, http://genetics.cs.ucla.edu/meta; Matrix eQTL software, http://www.bios.unc.edu/research/genomic_software/Matrix_eQTL.

## Acknowledgements

This work was supported by NIH grants MH090951 and HG02585 to M.S., and by a grant from the Gordon and Betty Moore Foundation (GBMF #4559) to M.S. S.M.U. was supported by NIH grant T32HD007009. Expression (GTEx) Project was supported by the Common Fund of the Office of the Director of the National Institutes of Health. Additional funds were provided by the NCI, NHGRI, NHLBI, NIDA, NIMH, and NINDS. Donors were enrolled at Biospecimen Source Sites funded by NCI/SAIC-Frederick, Inc. (SAIC-F) subcontracts to the National Disease Research Interchange (10XS170), Roswell Park Cancer Institute (10XS171), and Science Care, Inc. (X10S172). The Laboratory, Data Analysis, and Coordinating Center (LDACC) was funded through a contract (HHSN268201000029C) to The Broad Institute, Inc. Biorepository operations were funded through an SAIC-F subcontract to Van Andel Institute (10ST1035). Additional data repository and project management were provided by SAIC-F (HHSN261200800001E). The Brain Bank was supported by a supplement to University of Miami grants DA006227 & DA033684 and to contract N01MH000028. Statistical Methods development grants were made to the University of Geneva (MH090941 & MH101814), the University of Chicago (MH090951, MH090937, MH101820, MH101825), the University of North Carolina-Chapel Hill (MH090936 & MH101819), Harvard University (MH090948), Stanford University (MH101782), Washington University St. Louis (MH101810), and the University of Pennsylvania (MH101822). The data used for the analyses described in this manuscript were obtained from the GTEx Portal on October 17, 2015.

## Author contributions

S.M.U. and M.S. conceived of the project and developed the statistical methods. S.M.U. implemented the comparisons with simulated data. S.M.U. and G.W. performed the analyses of the GTEx data, and additional analyses. S.M.U., G.W. and M.S. implemented the software, with contributions from P.C. S.M.U. and M.S. wrote the manuscript, with input from G.W. and P.C. P.C. and G.W. prepared the online code and data resources.

## Competing interests The authors declare no competing financial interests

## Methods

### Model and fitting

Let *b*_*ir*_ (*j* = 1,…, *J*; *r* = 1,…, *R*) denote the true value of effect *j* in condition *r*.

Further, let *b*̂_*jr*_ denote the “observed” estimate of this effect, and let *s*̂_*jr*_ be the standard error of this estimate, so *z*_*jr*_ = *b*̂_*jr*_/*s*̂_*jr*_ is the standard *Z* statistic used to test whether *b*_*jr*_ is zero. Let *B*, *B*̂, *S* and *Z* denote the corresponding *J* × *R* matrices, and let *b*_*j*_ (respectively, *b*̂_*j*_, *z*_*j*_) denote the *j*th row of *B* (respectively, *B*̂, *Z*).

We assume *b*̂_*j*_ is normally distributed about *b*_*j*_, with variance-covariance matrix *V*_*j*_ (defined below), and that *b*_*j*_ follows eq. 1:

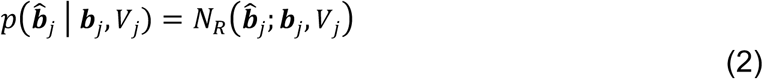

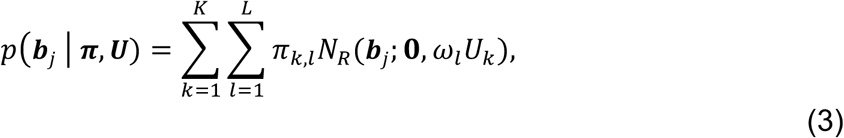

where *N*_*R*_(·; *μ*, Σ) denotes the multivariate normal (MVN) density in *R* dimensions with mean *μ* and variance-covariance matrix Σ. Here, each *U*_*k*_ is a covariance matrix that captures a pattern of effects; each *ω*_*l*_. is a scaling coefficient that corresponds to a different effect size; and the mixture proportions *π*_*k*,*l*_ determine the relative frequency of each covariance-scale combination. The scaling coefficients *ω*_*l*_. take values on a fixed dense grid that spans “very small” to “very large” so as to capture the full range of effects that could occur.

Combining eqs. 2 and 3 implies that the marginal distribution of *b*̂_*j*_, integrating out *b*_*j*_, is

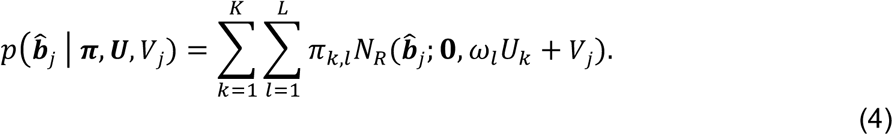

Each covariance matrix *V*_*j*_ is specified as *V*_*j*_ = *S*_*j*_*CS*_*j*_, where *C* is a correlation matrix that accounts for correlations among the measurements in the *R* conditions, and *S*_*j*_ is the *R* × *R* diagonal matrix with diagonal elements *s*̂_*j*1_, …, *s*̂_*jR*_. If measurements in the *R* conditions are independent, one would set *C* = *I*_*R*_, the *R* × *R* identity matrix, and therefore 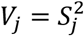. However, in our GTEx analysis the measurements are correlated due to sample overlap (individuals in common) among tissues; we estimate this correlation from the data (see “Estimating the correlation matrix *C*” below). The methods place no restriction on the *V*_*j*_ except that they must be symmetric positive definite matrices.

The steps of `mash` are:

1. Estimate *U* and *π*. This involves two sub-steps:

a. Define a set of (normalized) data-driven and canonical covariance matrices, *Û*.
b. Given *Û*, estimate *π* by maximum likelihood. (A key idea is that if some matrices generated in Step 1a do not help capture patterns in the data, they will receive little weight.) Let *π*̂ denote this estimate.
2. Compute, for each *j*, the posterior distribution *p*(*b*_*j*_ | *b*̂_*j*_, *π*̂, *Û*, *V*_*j*_).

These steps are now detailed in turn.

### Generate data-driven covariance matrices *U*_*k*_

We first identify rows *j* of matrix *B*̂ that correspond to the “strongest” effects. For example, in the GTEx data we chose rows corresponding to the “top” SNP for each gene, which we defined to be the SNP with the highest value of 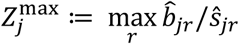. (We used the maximum rather than the sum because we wanted to include effects that were strong in a single condition, not just effects that were shared among conditions.) For the simulated data, we ran ash separately for each condition *r*, computed *lfsr*_*jr*_ for each (*j*, *r*), then chose rows *j* for which 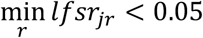.

Next, we fit a mixture of MVN distributions to these strongest effects using methods from Bovy *et al*.^35^. Bovy *et al*. describe an EM algorithm for fitting a model similar to eqs. 2 and 3, with the crucial difference that there are no scaling parameters on the covariances:

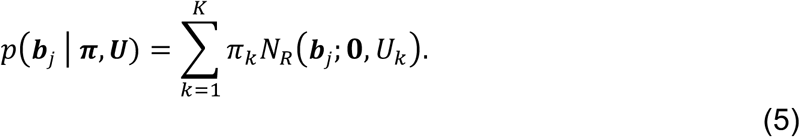

The absence of the scaling factors *ω*_*l*_. means that, compared with `mash`, their model is less well suited to capturing effects that have similar patterns (relative sizes across conditions) but vary in magnitude. By applying it to only the largest effects, we seek to sidestep this issue.

Estimates of *U*_*k*_ from this EM algorithm are sensitive to initialization. Furthermore, we noticed an interesting feature of the EM algorithm: each iteration preserves the rank of the matrices *U*_*k*_, so that ranks of the estimated matrices are the same as the ranks of the matrices used to initialize the algorithm. We exploited this fact to ensure that some of the estimated *U*_*k*_ had low rank. This helped stabilize the estimates since rank-penalization is one way to regularize covariance matrix estimation.

To describe the initialization in detail, let *J*̃ denote the number of “strongest effects” selected above, and let *Z*̃ denote the column-centered *J*̃ × *R* matrix of *Z* scores for these “strong effects”. To extract the main patterns in *Z*̃, we perform dimension reduction on *Z*̃; specifically, we apply principal components analysis (through a singular value decomposition, or SVD) and sparse factor analysis^21^ (SFA) to *Z*̃. SVD yields a set of singular values and singular vectors of *Z*̃. Let *λ*_*p*_, *v*_*p*_ denote the *p*th singular value and corresponding right singular vector. SFA yields matrix factorization

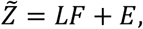

where *L* is a sparse *J* × *Q* matrix of loadings and *F* is a *Q* × *R* matrix of factors. We use *Q* = 5.

Given this matrix factorization, we use the EM algorithm to fit the mixture model (eq. 5) with *K* = 3, in which the three matrices *U*_*k*_ are initialized as follows:

- *U*_1_ = *Z*̃^*T*^*Z*̃/*J*̃, the empirical covariance matrix of *Z*̃.
- 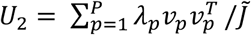, which is a rank-*P* approximation of the covariance matrix of *Z*̃, with *P* < *Q*. We use *P* = 3.
- *U*_3_ = *F*^*T*^*L*^*T*^*LF*/*J*̃, which is a rank-Q approximation of the covariance matrix of *Z*̃.

The output of the EM algorithm defines *U*1, *U*2 and *U*3 in the `mash` model (eq. 1).

In addition to the covariance matrices obtained from this EM algorithm, we define more covariance matrices from the SFA results; specifically, the *Q* = 5 rank-1 matrices 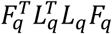, with *q* = 1,…, *Q*. The rationale for adding these rank-1 covariance matrices is that the SFA factors may directly reflect effect patterns in the data, and if so these matrices will be a helpful addition. (If they are not helpful, they will receive little weight when estimating *π*).

In summary, SFA and the EM algorithm generate 8 data-driven covariance matrices for our analysis of the GTEx and simulated data sets.

### Generate canonical covariance matrices *U*_*k*_

We use the following “canonical” covariance matrices:

- The identity matrix, *I*_*R*_. This represents the situation where the effects in different conditions are independent, which may be unlikely in some applications (like the GTEx application here), but seems useful to include if only to show that the data assign low weight to it.
- R rank-1 matrices 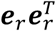, where *e*_*r*_ denotes the unit vector with zeros everywhere except for element *r*, which is set to 1. These matrices represent effects that occur in only one condition.
- The rank-1 matrix **11**^*T*^, where **1** denotes the R-vector of ones—this is the matrix of all ones, and represents effects that are identical among all conditions.

The user can, if desired, add other canonical matrices. For example, if *R* is moderate then one could add the 2^*R*-1^ canonical matrices that correspond to shared (equal) effects in each of the 2^*R*-1^ subsets.

In summary, this procedure produces 46 canonical covariance matrices for our analyses of the GTEx and simulated data sets, with *R* = 44.

### Standardize the covariance matrices

Since eq. 3 uses the same grid of scaling factors for each covariance matrix *U*_*k*_, we standardize the matrices *U*_*k*_ so that they are similar in scale. Specifically, for each *k*, we divide all elements *U*_*k*_ by the maximum diagonal element of *U*_*k*_ so that the maximum diagonal element of the rescaled matrix is 1. These rescaled matrices define *Û* in Step 1a of `mash`.

### Define grid of scaling factors

We define a dense grid of scaling factors, ranging from “very small” to “very large”. Stephens^16^ suggests a way to select suitable limits (*ω*_min_, *ω*_max_) for this grid in the univariate case; we apply this method to each condition *r*, and take the smallest *ω*_min_ and the largest *ω*_max_ as the grid limits. The internal points of the grid are obtained, as in the univariate case^16^, by setting *ω*_*l*_ = *ω*_max_/*m*^*l*-1^, for *l* = 1,…,*L*, where *m* > 1 is a user-tunable parameter that controls the grid density, and *L* is chosen to be just large enough so that *ω*_*L*_ < *ω*_min_. We use 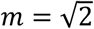.

### Estimate *π* by maximum likelihood

Given *Û* and *ω*, we estimate the mixture proportions *π* by maximum likelihood.

To simplify notation, let Σ_*k*, *l*_ := *ω*_*l*_*U*_*k*_, and replace the double index *k*, *l* with a single index *p* ranging from 1 to *P* := *KL*, so that eqs. 3 & 4 become

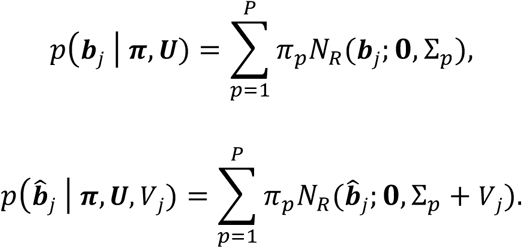

Assuming independence of the rows of *B*̂, the likelihood for *π* is

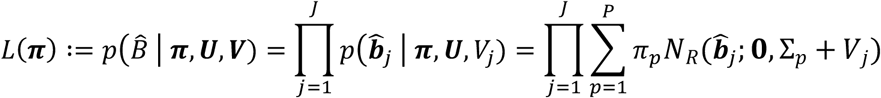

If the rows of *B*̂ are not independent, this may be interpreted as a “composite likelihood”^36^, which generally yields consistent point estimates. (We also empirically confirmed that our GTEx results were robust to LD pruning of SNPs; see Supplementary Fig. 9.)

Maximizing L(*π*) is a convex optimization problem, which we solve using EM^37^, accelerated using SQUAREM^38^. If *B*̂ has a large number of rows, we can reduce computational effort by taking a random subset of rows. In the GTEx analysis, we use a random subset of 20,000 rows. (It is important that this is a random subset, and not just the *J*̃ rows of strong effects.)

### Posterior calculations

If *b* ~ *N*_*R*_(0,*U*) and *b*̂|*b* ~ *N*_*R*_(*b*, *V*), then

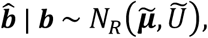

where

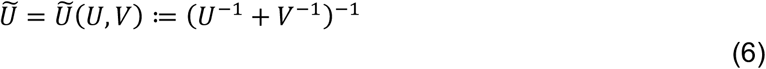

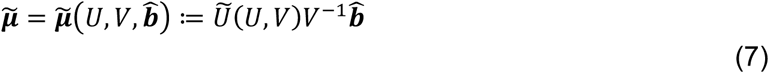

This result is easily extended to the case where the prior on *b* is a mixture of MVNs (eq. 3). In this case the posterior distribution is

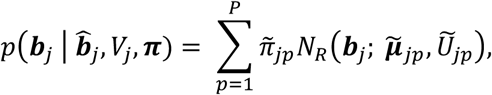

where *U*̃_*ip*_ = *U*̃(Σ_*p*_, *V*_*j*_) (eq. 6) and *μ*̃ := *μ*̃(Σ_*p*_, *V*_*j*_, *b*̂_*j*_) (eq. 7), and

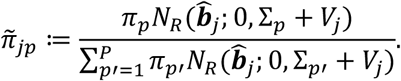

The posterior mean and variance are

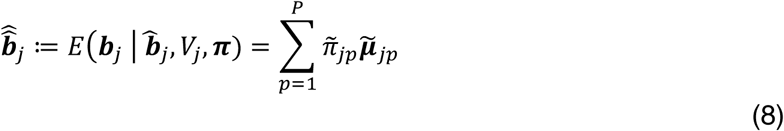

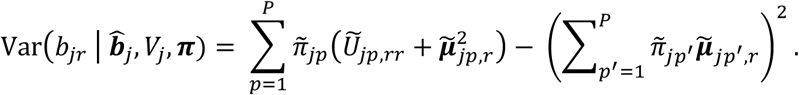

The local false sign rate (*lfsr*) is defined by Stephens^16^ as:

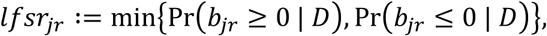

where *D* denotes the available data. Intuitively, *lfsr*_*jr*_ is the probability that we would incorrectly predict the sign of the effect if we were to use our best guess of the sign (positive or negative). Thus, a small *lfsr* indicates high confidence in the sign of an effect. The *lfsr* is more conservative than the local false discovery rate (*lfdr*)^39^ because requiring high confidence in the sign is more stringent than requiring high confidence that the effect be non-zero. More importantly, the *lfsr* is more robust to modelling assumptions than the *lfdr*^16^, a particularly important issue in multivariate analyses where modelling assumptions play a larger role.

### Bayes Factors testing global null

The Bayes Factor for *b* ≠ 0 against the null model (*b* = 0) is

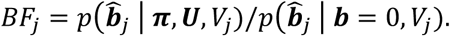

The numerator is given by eq. 4, and the denominator is given by eq. 2, with *b* = 0.

### The EZ model, and applying mash to *Z* scores

The model (eq. 3) assumes the *b*_*j*_ are independent of their standard errors, *V*_*j*_. We refer to this as the “Exchangeable Effects” (EE) model^23^. An alternative is to allow the effects to scale with standard error, so that effects with larger standard error tend to be larger,

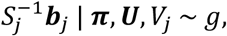

where *g* is shorthand for the mixture of MVNs (eq. 3). We refer to this as the “Exchangeable *Z*” (EZ) model because the left of this equation is the vector of *Z* scores for effect *j*.

As explained in Stephens^16^, the EZ model can be fit by applying the same code as the EE model to the *Z* statistics instad of the effect estimates, *B*̂, in which the standard errors of the *Z* statistics are set to 1; in other words, set *b*̂_*jr*_ = *z*_*jr*_ and *s*̂_*jr*_ = 1. (The correlation matrix, *C*, remains the same.) One advantage of the EZ model is that it can be fit using only the *Z* scores; it does not require access to both the effect estimates and their standard errors. The *lfsr* can also be computed using only the *Z* scores. However, the posterior mean estimates that arise from this model are estimates of 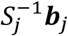; transforming these to estimates of effect sizes *b*_*j*_ requires knowledge of *S*_*j*_.

We analyzed the GTEx data using both EE and EZ models. Results were qualitatively similar in terms of patterns of sharing, but the EZ model performed better in cross-validation tests of model fit (see below), so we used results from the EZ model.

### Estimating the correlation matrix *C*

*C* is an *R* × *R* correlation matrix that accounts for correlations among measurements in the *R* conditions. To estimate this matrix, we exploit the fact that *C* is the correlation matrix of the *Z* scores under the null (*b*_*j*_ = 0). Specifically, we estimate *C* from the empirical correlation matrix of the *Z* scores for the effects *j* that are most consistent with the null,

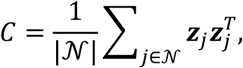

where 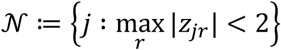.

For the GTEx data, we found that measurements in different tissues were not highly correlated; all elements of estimated *C* had magnitude less than 0.2, and 95% had magnitude less than 0.1. However, in cross-validation tests (below), this estimated *C* produced better model fit than ignoring correlations (*i.e., C* = *I*_*R*_).

### Cross-validation of model fit

To compare model fitting procedures, we randomly divide the rows of the data matrix *B*̂ into two subsets—a “training set” *B*̂_*train*_, and a “test set” *B*̂_*test*_. We then apply `mash` to *B*̂_*train*_, yielding estimates (*π*̂, *Û*), and assess the “fit” of these estimates by computing the log-likelihood in the test data, log *p*(*B*̂_*test*_|*π*̂, *Û*, *V*), which is given by eq. 4. This cross-validation strategy can also be used to compare different approaches to estimating *Û*, and our current strategy was developed and refined using this framework.

When applying cross-validation to the GTEx data, we create test and training sets by randomly selecting half the genes, rather than half the rows (gene-SNP pairs). Specifically, we use genes on even-numbered chromosomes as the training set, and genes on odd-numbered chromosomes as the test set. This ensures that rows in the test set were independent of rows in the training set.

### Visualizing *U*_*k*_

In the simulations, and in our application to the GTEx data, *R* = 44, so each *U*_*k*_ is a 44 × 44 covariance matrix, and each component of the mixture (eq. 1) is a distribution in 44 dimensions. Visualizing such a distribution is challenging. However, we can get some insight from the first eigenvector of *U*_*k*_, denoted by *v*_*k*_, which captures the principal direction of the effects. If *U*_*k*_ is dominated by this principal direction, then we can think of effects from that component as being of the form *Xv*_*k*_ for some scalar *λ*. For example, if the elements of the vector *v*_*k*_ are approximately equal, then component *k* captures effects that are approximately equal in all conditions; alternatively, if *v*_*k*_ has one large element, with other elements close to zero, then component *k* corresponds to an effect that is strong in only one condition. See Fig. 3 and Supplementary Fig. 2 for illustration.

### RRMSE (accuracy of estimates in simulation studies)

In simulations, we used the “relative root mean squared error” (RRMSE) to assess error in effect estimates (e.g., Fig. 2). Let 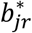 denote the estimate of effect *j* in condition *r* (for `mash`, 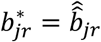). Then

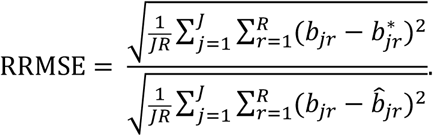

### ROC curves

For the ROC curves in Fig. 2c–d, the True Positive Rate (TPR) and False Positive Rate (FPR) are computed at any given threshold *t* as

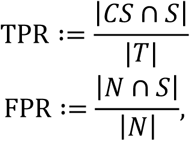

where *S* is the set of significant results at threshold *t, CS* the set of correctly signed results, *T* the set of true (non-zero) effects and *N* the set of null effects:

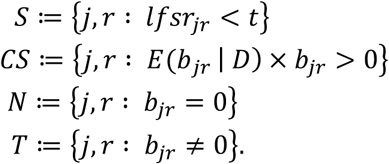

Thus, to be considered a true positive, we require that an effect be both “significant” and correctly signed.

For the ROC curves in Fig. 2a–b, the TPR and FPR are computed by treating units *j* as “discoveries”. For example, suppose a method produces a *p* value *p*_*j*_ for testing unit *j*. Then, at any threshold *t*, the TPR and FPR are

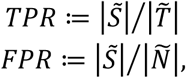

where *S*̃ is the set of significant units at threshold *t*, *T*̃ the set of true (non-null) units, and *N*̃ the set of null units:

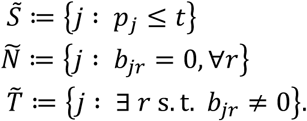

### Effective sample size

We define the “effective sample size” (ESS) for tissue *r* to be

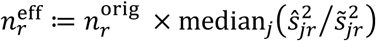

where 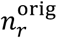 is the number of available samples in tissue *r* (Supplementary Table 3), and *s*̂_*jr*_ and *s*̃_*jr*_ are the standard error and posterior standard deviation, respectively, for effect *j* in tissue *r*.

### Normalized effects

For estimating sharing of effects by magnitude (e.g., Supplementary Fig. 7), we define a “normalized” effect *b*̃ in each condition as the ratio of its effect in that condition to the largest effect across all conditions:

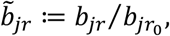

where

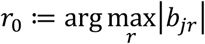

For example, in the application to multi-tissue eQTL analysis, a normalized effect *b*̃_*jr*_ = 0.5 means that the effect of eQTL *j* in tissue *r* is half that of its effect in the strongest tissue.

### Sharing by sign and magnitude

We define two effects to be “shared in sign” if they have the same sign; we define them to be “shared in magnitude” if they *also* have an effect within a factor of 2 of one another. (In some applications it may make sense to drop the requirement that effects are of the same sign when assessing sharing by magnitude, but not here.)

In Fig. 5 and Table 1, we assess sharing for each eQTL by comparing its estimated effect in each tissue against the strongest estimated effect across all tissues. Thus, for example, if an eQTL has a very strong positive effect in one tissue and weak negative effects in all other tissues, then the number of tissues “shared by sign” for this eQTL would be 1.

In Fig. 6 (and Supplementary Fig. 5), we investigate sharing between each pair of tissues.

When computing sharing between tissues *r* and *s*, we consider only QTLs that are “significant” (*lfsr* < 0.05) in at least one of *r* and *s*. This is to avoid sharing estimates being driven by estimates of null or nearly-null effects.

More formally, define

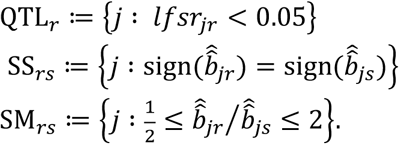

Then the sharing by sign between *r* and *s* is given by

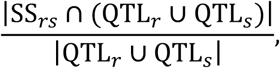

and sharing by magnitude between *r* and *s* is given by

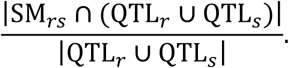

These estimates of sharing are based on the posterior estimates 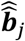, to simplify their calculation.

However, we obtained similar estimates of sharing when we accounted for uncertainty in 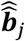 by sampling from the posterior distributions of the effects.

`ash`: We applied ash^16^ separately to each tissue using the same 20,000 randomly-selected gene-SNP pairs as for `mash`. We then computed the posterior means and *lfsr* for the top SNPs.

`mash-bmalite`: This is a simplification of `mash` without the data-driven covariance matrices, and with particular choices for the canonical covariance matrices and scaling factors. It can be thought of as a version of `BMAlite`^5^ that outputs effect size estimates and *lfsr* values. (The original software does not compute these quantities.)

Specifically, the *U* for `mash-bmalite` includes the 44 singleton configurations 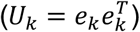, and matrices corresponding to the models in Flutre *et al*.^5^, with heterogeneity parameters *H* = {0, 0. 25, 0.5, 1}. (When *H* = 0, effects are equal in all conditions; when *H* = 1, effects are independent among conditions.) We used *ω* = {0.1, 0.4, 1.6, 6.4, 25.6}, similar to the coarse grid used in Flutre *et al*.^5^, and chosen to capture the full range of the GTEx (real and simulated) *Z* statistics.

### Statistical tests

For the ash, `mash` and `mash-bmalite` methods, an association was considered significant if the *lfsr*^16^ was less than 0.05. For all metasoft approaches included in the simulation experiments (FE, RE2, BE), *p* values were computed as described in ^9,14^. The metasoft *p* value threshold was varied to assess performance in the simulation experiments (Fig. 2).

### “Shared, structured effects” simulations

We simulated each *b*_*j*_ from the mixture model (eq. 3), with mixture parameters based on fit to the GTEx data. This is the simulation procedure in more detail:

1. We took the 8 “data-driven” covariance matrices *U*_1_,…, *U*_8_ learned from the GTEx data, standardized as described above.
2. We simulated 400 “non-null units”: independently for *j* = 1, …, 400, we (a) chose a component *k* uniformly at random from 1,…, 8, (b) simulated a scaling factor *ω* as the absolute value of an *N*(0,1) random variable, and (c) simulated *b*_*j*_ ~ *N*_44_(0, *ωU*_*k*_).
3. For the 19,600 “null units”, we set *b*_*j*_ = 0.
4. For all 20,000 units, we simulated *b*̂_*j*_ ~ *N*_44_(*b*_*j*_, *V*_*j*_), where *V*_*j*_ was the diagonal matrix with diagonal elements 0.1^2^ (so that standard errors were 0.1, similar to the estimates from the GTEx data).

### “Shared, unstructured effects” simulations

We simulated as for “shared, structured effects”, except that the 400 non-null units in Step 2c were simulated as *b*_*j*_ ~ *N*_44_(**0**,0.1^2^*I*).

### “Independent effects” simulations

We also simulated data from effects that were entirely independent across conditions:

1. Independently for each *r* = 1, …, 44, we chose a random set of 400 *j* ∈ {1,…, 2 × 10^4^} to be the “true effects”.
2. For the “true effects”, we simulated *b*_*jr*_ ~ *N*(0,Σ^2^), where Σ^2^ was chosen with equal probability from the set {0.1, 0.5, 0.75, 1}. In this way, the true effects captured both small and large effects for each condition. All other effects were set to zero.
3. We simulated *b*_*j*_ ~ *N*_44_(*b*_*j*_, *V*_*j*_) as in the other simulations.

### Analysis of simulated data

Each simulated dataset was analyzed using `mash`, following the procedure described above. Note that we re-estimated (*U*, *π*) from the data without making use of the true values used to simulate the data. We estimated effects by their posterior means (eq. 8), and assessed significance by the *lfsr*. The ash and `mash`-bmalite methods were applied to the simulated data in the same way that they were applied to the GTEx data (see above).

### Data availability

The GTEx study data are available through dbGaP under accession phs000424.v6.p1. The GTEx summary statistics used in the `mash` analysis have been deposited in Zenodo (doi:10.5281/zenodo.1296399).

### Code availability

Source code implementing our methods, including processing and analysis of the GTEx data, adaptive shrinkage (“ash”) and multivariate adaptive shrinkage (“mash”, “mash-bmalite”), is available online, and is distributed under GNU and BSD open source licenses (see URLs).

## Supplementary Note

### Multivariate adaptive shrinkage (mash), and its relationship with existing methods

An important novel feature of our method, `mash`, is its focus on *estimation of effect sizes*, in contrast to most existing multivariate analysis methods that focus only on *testing* for non-zero effects. Further, *mash* is more than just an extension of existing methods to estimate effect sizes because the underlying model is more flexible than models underlying existing methods—and, indeed, includes existing models as special cases.

The `mash` method includes many existing methods for joint analysis of multiple effects as special cases. Specifically, many existing methods correspond to making particular choices for the set of “canonical” covariance matrices (with no data-driven matrices). For example, a simple “fixed effects” meta-analysis—which assumes equal effects in all conditions—corresponds to *K* = 1 with *U*_1_ = 11^*T*^ (a matrix of all ones). (This covariance matrix is singular, but it is still allowed in `mash`.) A more flexible assumption is that effects in different conditions are normally distributed about some mean—this corresponds to the multivariate normal assumption made in `mash` if the mean is assumed to be normally distributed as in Wen & Stephens^1^. More flexible still are models that allow effects to be exactly zero in subsets of conditions, as in Flutre *et al*.^2^ and Li *et al*.^3^. These models correspond to using (singular) covariances *Uk* with zeros in the rows and columns corresponding to the subset of conditions with no effect.

`mash` extends the capabilities of previous methods in two ways: first, `mash` includes a large number of scaling coefficients *ω*, which allows `mash` to flexibly capture a range of effect distributions^4^; second, and perhaps more importantly, `mash` includes data-driven covariance matrices, making it more flexible and adaptive to patterns in the data. This innovation is particularly helpful in settings with moderately large *R*—as in the GTEx data, with *R* = 44—where it becomes impractical to pre-specify canonical matrices for all patterns of sharing that might occur. For example, Flutre *et al*.^2^ and Li *et al*.^3^ consider all 2^*R*^ combinations of sparsity in the effects, which is feasible for *R* = 9 (see Flutre *et al*.^2^), but impractical at *R* = 44. While it is possible to restrict the number of combinations considered (e.g., BMAlite^2^), this comes at a cost to flexibility. The addition of data-driven covariance matrices helps to address this issue, making `mash` both flexible and computationally tractable for moderately large *R*.

In addition to effect estimates, `mash` also provides a measure of significance for each effect in each condition. Specifically, `mash` estimates the “local false sign rate” (*lfsr*)^4^, which is the probability that the effet is estimated with the incorrect sign. The *lfsr* is analogous to the local false discovery rate^6^, but is more stringent in that it insists that effects be correctly signed to be considered “true discoveries”. Similarly, `mash-bmalite` can estimate the *lfsr* (under its less flexible model), and `ash` can estimate the *lfsr* separately for each condition.

### Comparison with metasoft

Among existing software packages for this problem, `metasoft`^7^ is in some respects the most comparable to `mash`. In particular, it is both generic—requiring only effect estimates and their standard errors—and computationally tractable for *R* = 44. The `metasoft` software implements several different multivariate tests for association analysis, each corresponding to different multivariate models for the effects. For example, the FE model assumes that the effects in all conditions are equal; the RE2 model assumes that the effects are normally distributed about some common mean, with deviations from that mean being independent among conditions^8^; and the BE model is an extension of the RE2 model allowing that some effects are exactly zero^7^. These models are similar to the BMAlite models from Flutre *et al*.^2^, and none capture the kinds of structured effects that can be learned from the data by `mash`. However, because differences in software implementation sometimes lead to unanticipated differences in performance, we also performed simple direct benchmarks comparing `mash` and `mash-bmalite` with `metasoft`. For each model (FE, RE2, BE), `metasoft` produces a *p* value for each multivariate test, whereas `mash` and `mash-bmalite` produced a Bayes Factor (see Online Methods); in each case, these can be used to rank the significance of the tests.

### Assessing heterogeneity and sharing in effects

In analyses of effects in multiple conditions, it is often desirable to identify effects that are shared across many conditions, or, conversely, those that are specific to one or a few conditions. This is a particularly delicate task. For example, Flutre *et al*.^2^ emphasize that the simplest approach—first identifying significant signals separately in each condition, then examining the overlap of significant effects—can substantially underestimate sharing. This is due to incomplete power; by chance, a shared effect can easily be significant in one condition and not in another. To address this, Flutre *et al*.^2^ and Li *et al*.^3^ estimated sharing among conditions as a parameter in a joint hierarchical model, which takes account of incomplete power. However, these approaches are infeasible for *R* = 44. Furthermore, even for smaller values of *R* they have some drawbacks. In particular, they are based on a “binary” notion of sharing—*i.e*., whether an effect is non-zero in each condition—so they do not capture differences in magnitude or directions of effects among conditions. If effects shared among conditions differ greatly in magnitude—for example, being very strong in one condition and weak in all others—then this would seem useful to know.

We addressed this limitation by taking a new approach to quantify similarity of effects. Specifically, we assessed sharing of effects in two ways: (i) “sharing by sign” (estimates have the same direction); and (ii) “sharing by magnitude” (effects are similar in magnitude). We defined “similar in magnitude” to mean both the same sign and within a factor of 2 of one another. (Other thresholds could be used, and in some settings—e.g., when “conditions” are different phenotypes—the requirement that effects have the same sign could be dropped.) These measures of sharing can be computed for any pair of conditions, and an overall summary of sharing across conditions can be obtained by assessing how many conditions share with some reference condition. (We used the condition with the largest estimated effect as reference.) These measures of sharing could be naively estimated from the original effect estimates in each condition; however, errors in these effect estimates will naturally lead to errors in assessed sharing. Because `mash` combines information across conditions to improve effect estimates, it can also provide more accurate estimates of sharing.

### Effects of linkage disequilibrium

Linkage disequilibrium (LD) between SNPs has two distinct effects.

First, LD causes correlations in the observed effects of nearby SNPs for the same gene. This issue is likely to be minor here; `mash` ignores correlations between rows of *B*̂ when estimating the prior density g, and this can be justified as a “composite likelihood” approach^9^, which can perform well for computing joint estimates of model parameters.

Second, the effect estimates we obtained for each SNP from single-SNP analysis are not actually the individual causal effects of that SNP; rather, they are the *combined effects of all SNPs that are in LD with that SNP, weighted by their LD*^10,11^. This issue more likely has an impact because of the presence of multiple eQTLs in some or many genes. It also applies to all single-SNP eQTL analyses, which are the vast majority of all published eQTL analyses, and not just `mash`. Ideally, one would develop a multi-SNP, multi-tissue method for association analysis at each gene to avoid this issue. And, indeed, we see `mash` as a first step towards this more ambitious goal. However, for now we have limited this analysis to highlighting one specific feature of our results that we believe may be a consequence of the use of single-SNP effect estimates, and which will hopefully be better addressed as multi-SNP analyses are developed to better account for LD.

Specifically, we found that LD among multiple causal SNPs can cause single-SNP analyses to identify eQTLs that appear to have strong effects of opposite sign in different tissues. One example is shown in Supplementary Fig. 4; this eQTL has strong, positive *Z* scores in brain tissues, and negative *Z* scores in most other tissues. Initially, this suggested that this eQTL might have causal effects in opposite directions in brain versus non-brain tissues. However, there is another way to explain this result: there could be two eQTLs in LD with one another, one of which (e.g., eQTL A) has a strong effect in brain tissues, and the other of which (e.g., eQTL B) has a strong effect in other tissues. If the expression-increasing allele at eQTL A is in negative LD with the expression-increasing allele at eQTL B, then the single-SNP *Z* scores for both SNPs will show opposite signs in brain versus non-brain. Indeed, closer examination of the data at this particular gene suggests that this explanation is likely correct in this case (Supplementary Fig. 4). A similar example is discussed in the GTEx pilot study^5^ (their Supplementary Fig. 14).

Based on this reasoning, we believe that estimates of sharing in sign from single-SNP analyses such as ours are likely to be underestimates of the sharing in sign of actual causal effects. Therefore, we urge careful interpretation of an eQTL in multiple tissues that shows significant effects in different directions.

### Increase in effective sample size due to multivariate analysis

A feature of our multivariate analysis approach is improved quantitative estimates of effect sizes in each condition. When estimating effects in a single condition, `mash` uses the data not only from that condition but also from other “similar” conditions. In this way, `mash` effectively increases the available sample size, improving both accuracy and precision of estimates. The improvement will be greatest for conditions that are similar to many other conditions, and weakest for conditions with many “condition-specific” effects.

To illustrate this effect in the GTEx data, we computed an “effective sample size” (ESS) for each tissue based on the standard deviations of the `mash` estimates. The ESS estimates (Supplementary Fig. 6) vary from 240 for testis to 1,392 for coronary artery. Other tissues with smaller ESS include liver, pancreas, spleen and brain cerebellum. Identifying tissues with smaller ESS could help prioritize “under-represented” tissues in future experimental efforts.

For testis, the ESS of 240 represents only a small (1.4-fold) increase compared with actual sample size, reflecting that its effects are more “tissue specific”; that is, they are less correlated with other tissues. Other tissues showing a small gain in ESS include transformed fibroblasts and whole blood, which we also highlight for having more “tissue-specific” signals. By contrast, the ESS for coronary artery represents a 10-fold increase compared with the actual sample size for this tissue, reflecting its strong correlation with other tissues. On average across all tissues, `mash` provides a 4-fold increase in ESS for estimating the top eQTL effects, reflecting an overall moderate-to-large correlation in effect sizes across tissues.

One caveat of this analysis is that ESS reflects *average* gains in precision for a tissue; in practice, effects that are shared across many tissues will benefit more than effects that are tissue-specific. For example, if one were particularly interested in effects that are specific to uterus (which has the smallest actual sample size in our study), then the high reported ESS for uterus may not be as useful. In the end, detection of tissue-specific effects will benefit most from collecting more samples in the tissue of interest.

---

**Supplementary Figure 1.**
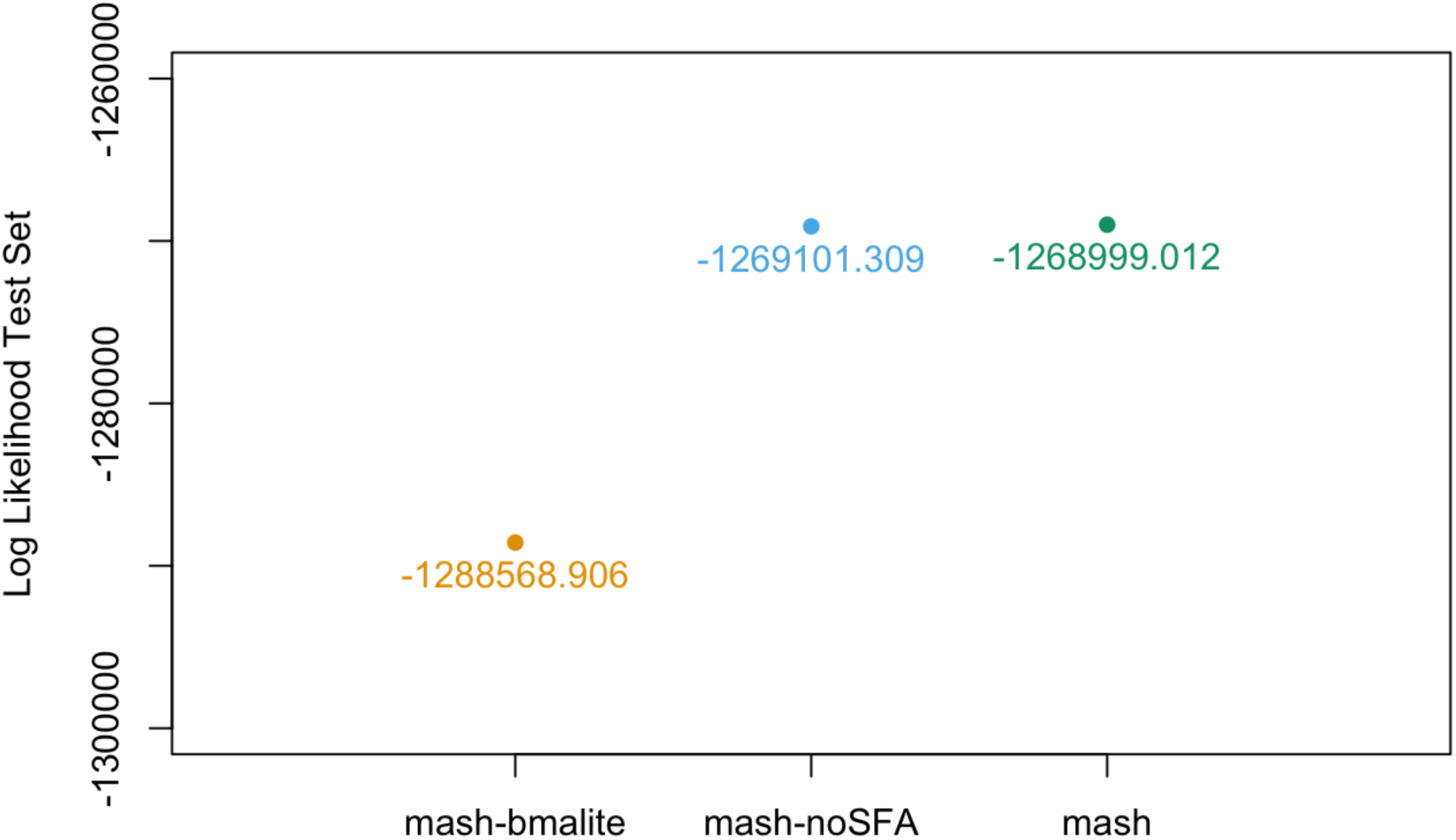
Increase in log-likelihood on test set as new *U*_*k*_ are added. The figure shows the log-likelihood on the test set for different “models” (choices of *U*_*k*_). From left to right, the models are: `mash-bmalite` (no data-driven *U*_*k*_); `mash-no-SFA` (the combination of canonical and data-driven covariances, excluding the rank-one matrices derived from SFA); `mash` (the full combination of canonical and data-driven covariances described here). The result illustrates how, as more data-driven covariances are added, the log-likelihood on the test set increases. Note that the difference in likelihood between the `mash` and `mash-no-SFA` is large—`mash` is approximately 100 log-likelihood units higher than `mash-no-SFA`—although this is difficult to see at this scale. Test-set log-likelihoods are based on *n* = 28,198 randomly selected gene-SNP pairs.

**Supplementary Figure 2.**
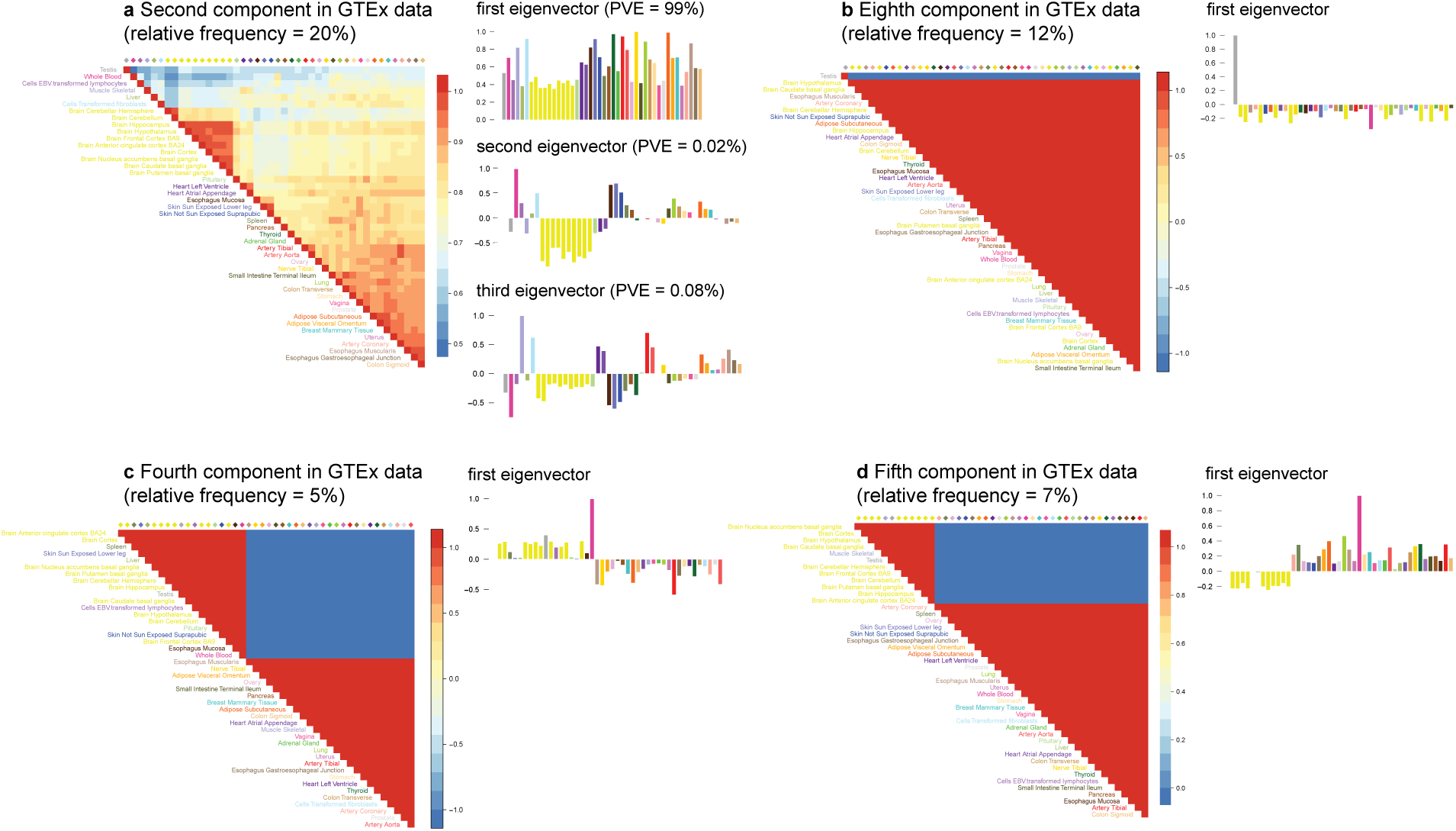
Summary of covariance matrices *U*_*k*_ with the largest estimated weights (>1%) in the GTEx data. For each covariance matrix *U*_*k*_, the figure shows the heatmap of the corresponding correlation matrix, and bar plots of the top eigenvectors of *U*_*k*_ (*n* = 16,069 independent gene-SNP pairs). Component 2 (**a**) captures qualitatively similar effects to the component shown in Fig. 3. Component 8 (**b**) captures testis-specific effects. Components 4 (**c**) and 5 (**d**) primarily capture effects that are stronger in whole blood than in other tissues.

**Supplementary Figure 3.**
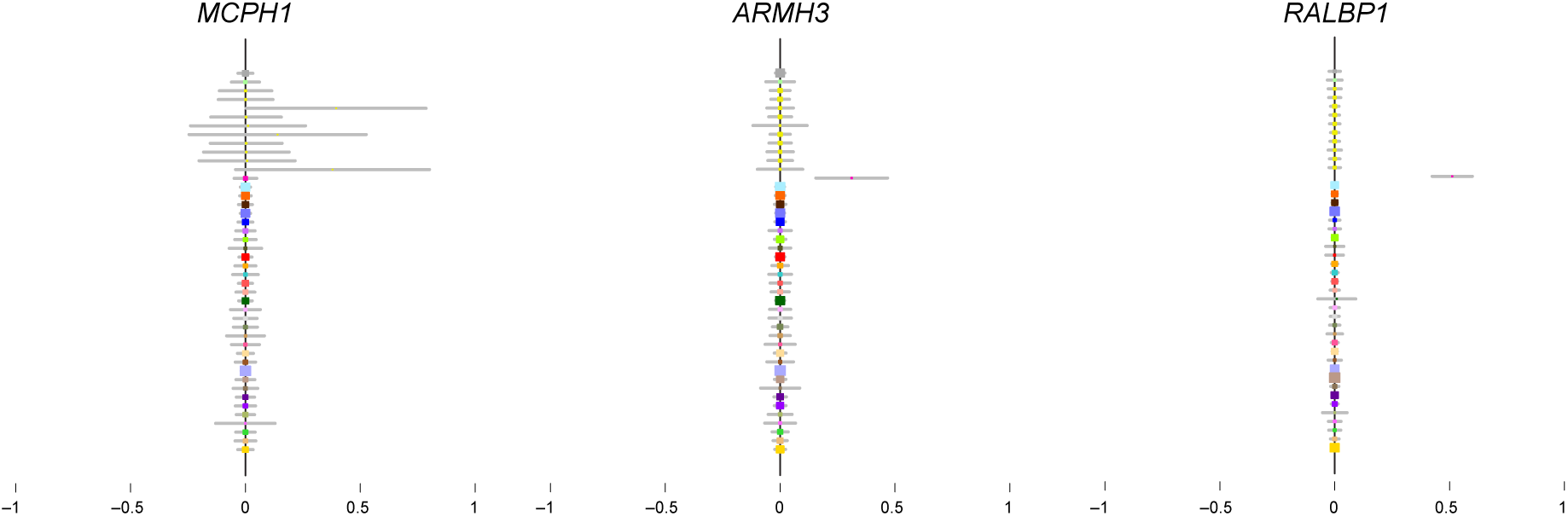
Estimates from the univariate method ash for the examples shown in Fig. 4. Each dot (color-coded as in Fig. 3) shows the effect estimate (posterior mean) from `ash`, with horizontal gray bars indicating ±2 posterior standard deviations. For all estimates, *n* = 83–430 individuals, depending on the tissue (Supplementary Table 3).

**Supplementary Figure 4.**
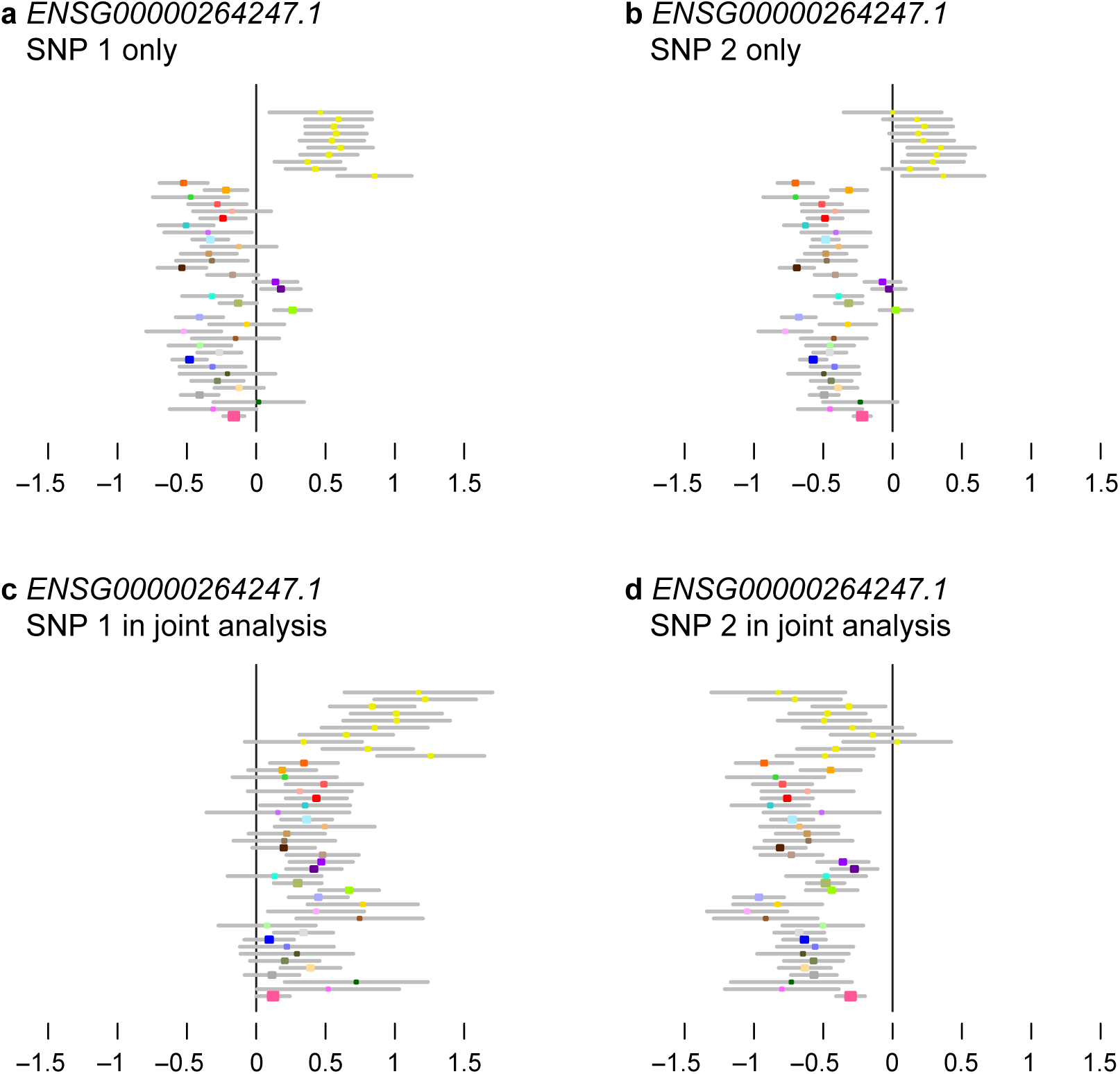
Illustration of how linkage disequilibrium (LD) can impact effect estimates. This gene was chosen as an example where the effect estimates in the “top eQTL” were opposite in sign in brain compared to non-brain tissues, and where further investigation suggested that this difference in effect directions could be explained by multiple eQTLs in LD. In this example, we define “SNP1” and “SNP2” as the SNPs that show the strongest eQTL associations in brain and non-brain tissues, respectively. The top panels show effect estimates for these SNPs from a simple (1-SNP) regression model in each tissue, *Y* = *μ* + *B*̂_*l*, *gi*_ where *i* in {1, 2} indexes the two SNPs. The bottom panels show effects from a multiple (2-SNP) regression model in each tissue, *Y* = *μ* + *B*̂_1_*g*_1_ + *B*̂_2_*g*_2_. Each dot shows the effect estimate for a single tissue (color-coded as in Fig. 3), with grey bars indicating ±2 standard errors. For all estimates, *n* = 83–430 individuals, depending on the tissue (Supplementary Table 3). The simple regression estimates (**a, b**) show opposite-direction effects in brain versus non-brain tissues (with testis and pituitary clustering with brain in one case). However, the multiple regression results (**c, d**) suggest that in fact there are (at least) two eQTLs in this gene, as SNP1 and SNP2 show a significant effect that excludes zero in most tissues. Furthermore, for both SNP1 and SNP2 the multiple regression effect estimates are consistent in sign across all tissues.

**Supplementary Figure 5.**
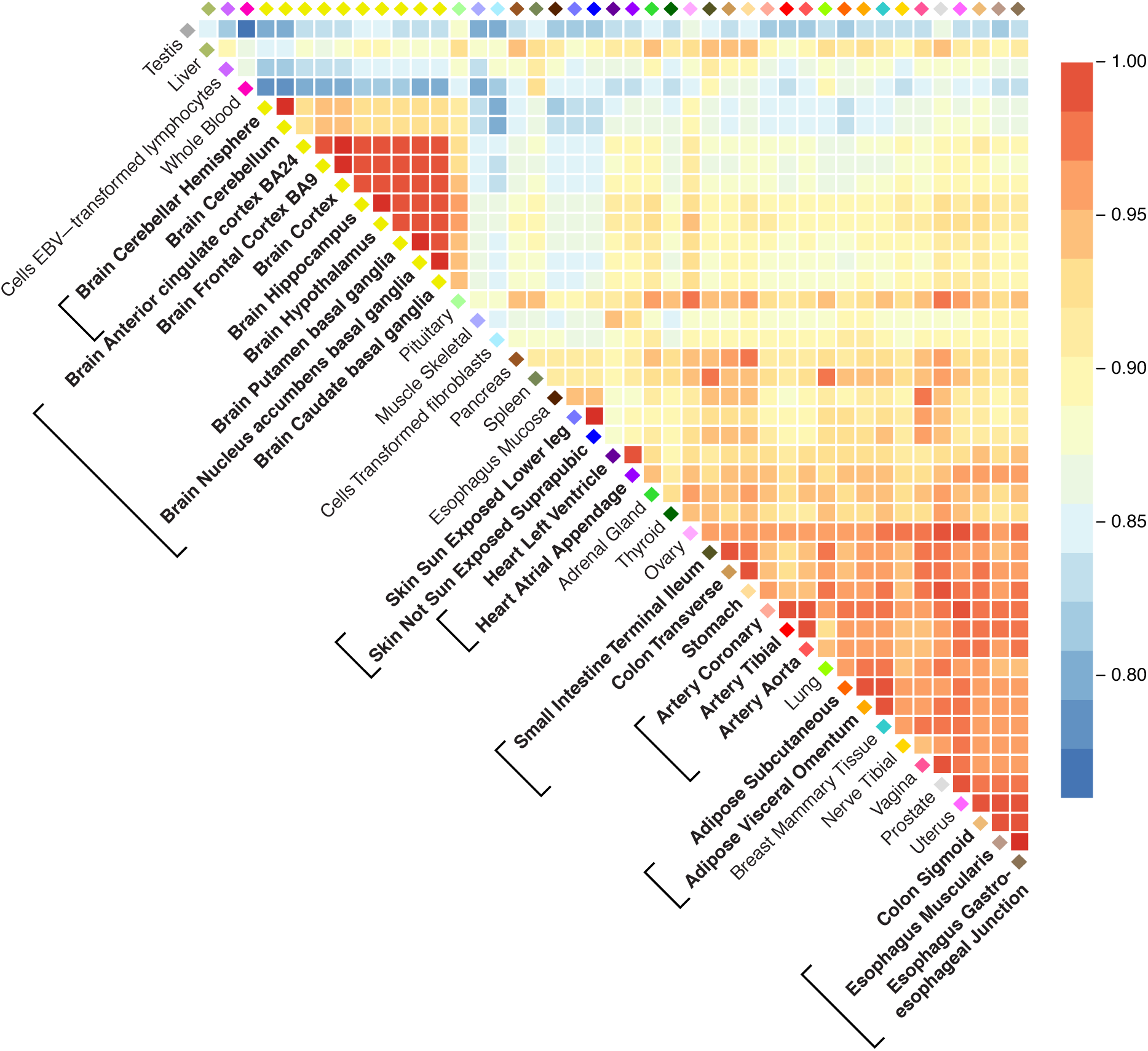
Pairwise sharing by sign. For each pair of tissues, we considered the top eQTLs that were significant (*lfsr* < 0.05) in at least one of the tissues, and calculated the proportion that have effect sizes with the same sign. These proportions are shown in this heatmap. *n* = 5,605–9,811 gene-SNP pairs, depending on pair of tissues compared.

**Supplementary Figure 6.**
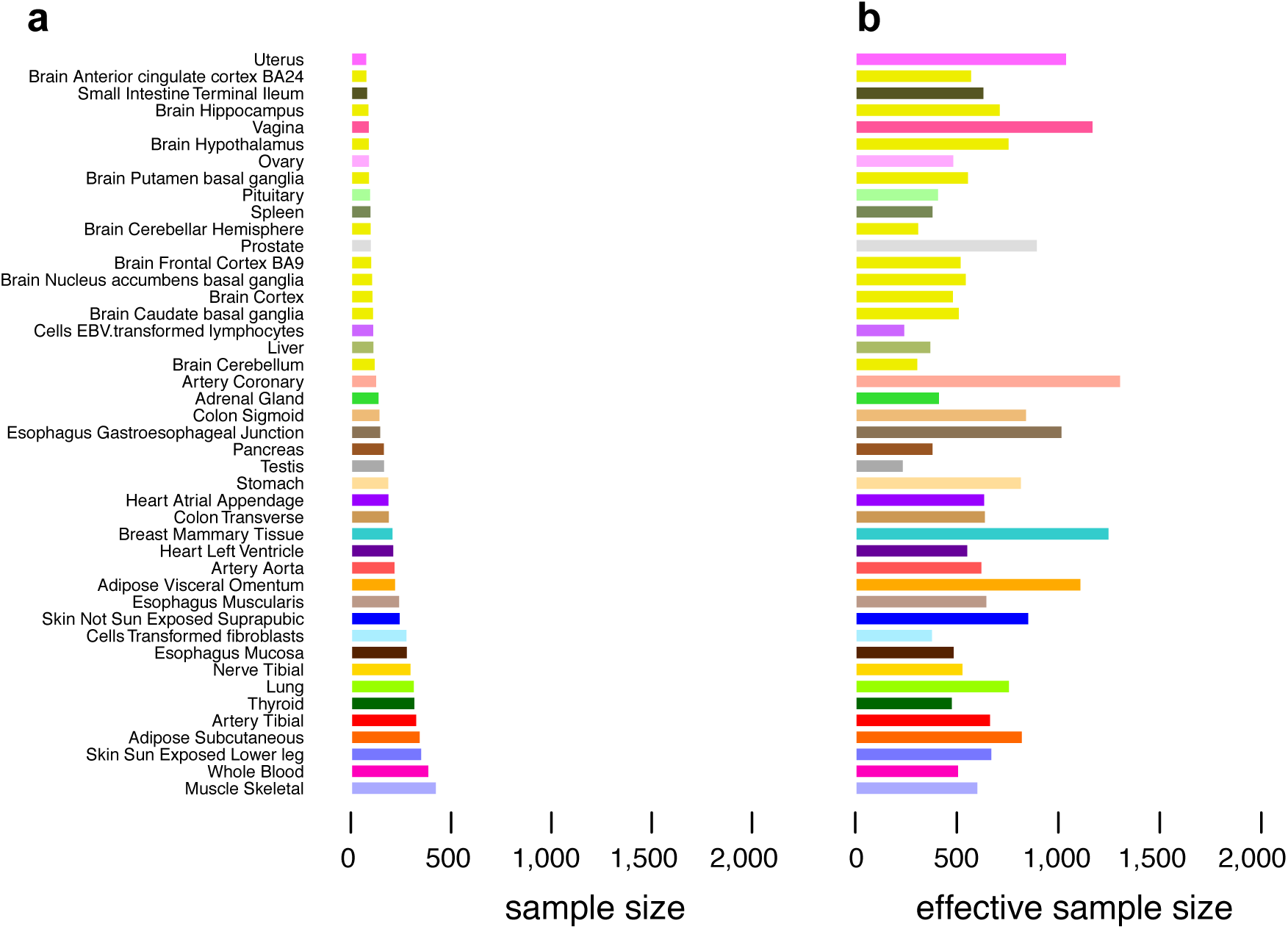
Sample sizes and effective sample sizes from mash analysis across tissues. Sample size (**a**) and median “effective sample size” (ESS) for each tissue (**b**). Tissues are ordered by their (original) sample size (Supplementary Table 3). Effective sample sizes are consistently higher than actual sample sizes, primarily due to sharing of information among tissues.

**Supplementary Figure 7.**
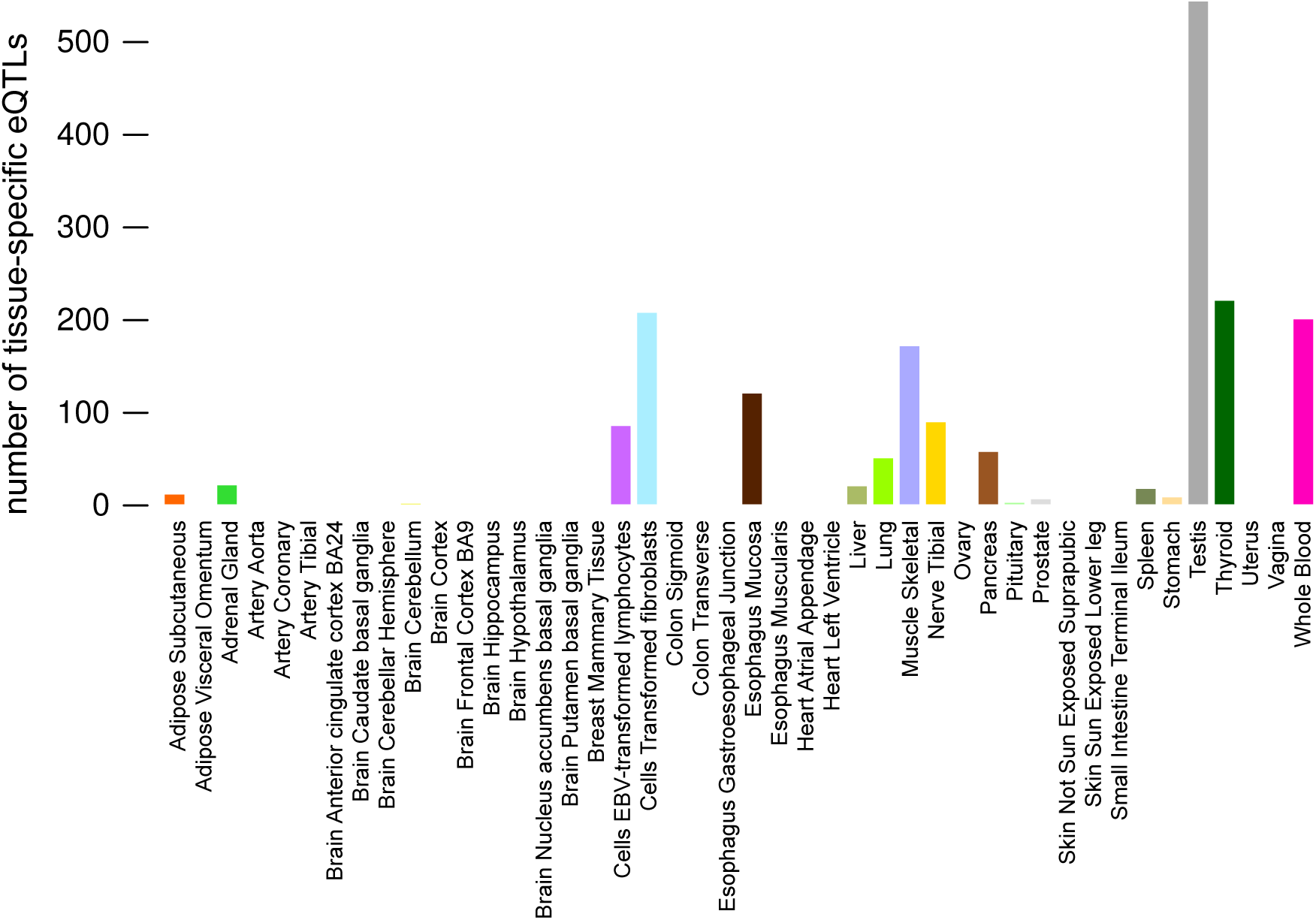
Number of tissue-specific eQTLs in each tissue. Here, “tissue-specific” is defined to mean that the effect is at least 2-fold larger in one tissue than in any other (*i.e*., *b*̃_*jr*_ > 0.5 in only one tissue).

**Supplementary Figure 8.**
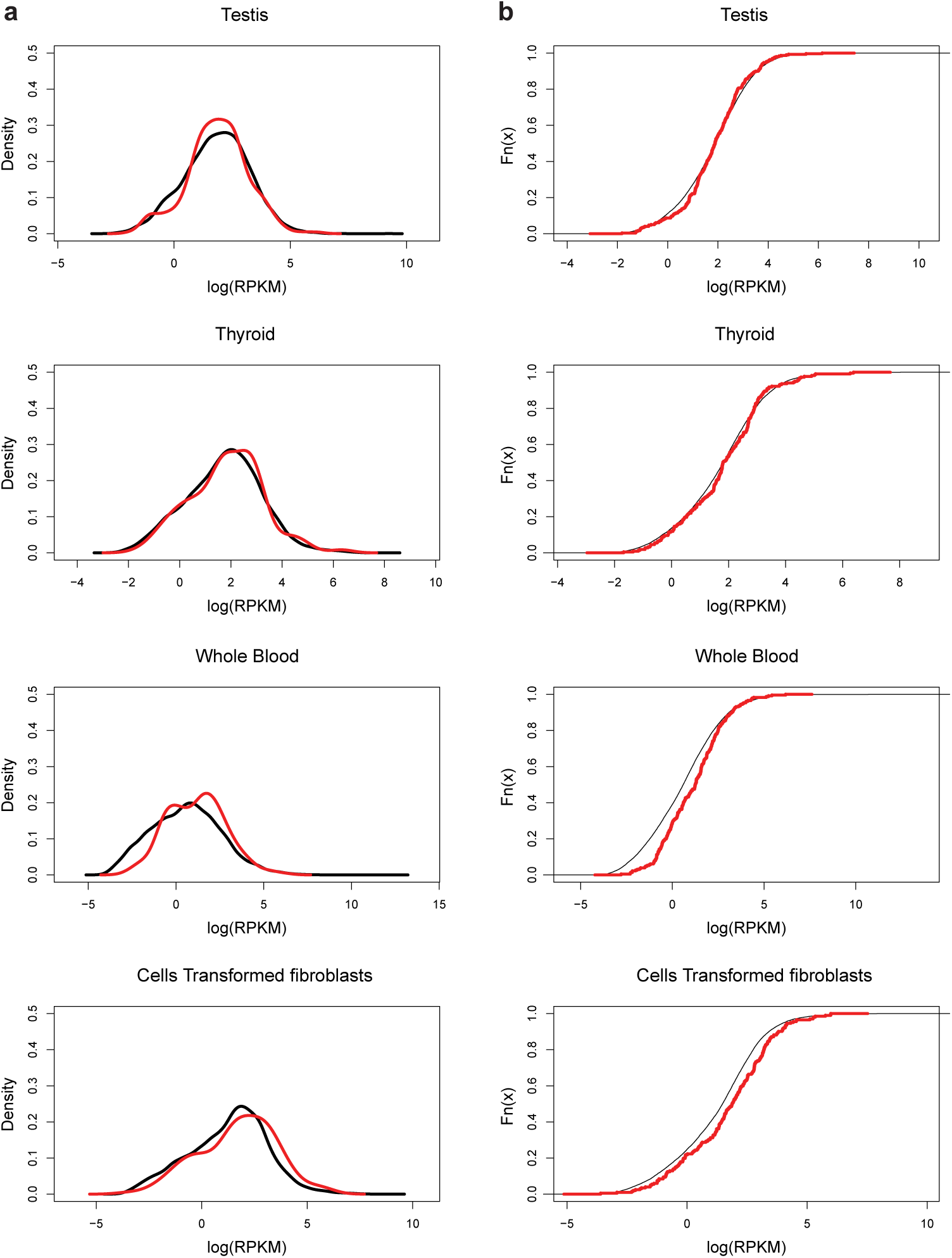
Expression levels in genes with tissue-specific eQTLs are similar to those in other genes. The plots compare the densities (**a**) and cumulative distribution functions (**b**) of the average expression level for genes identified as having a “tissue-specific” eQTL (red), and remaining genes (black), separately in four tissues—testis, thyroid, whole blood and transformed fibroblasts. In each case, the distribution functions are reasonably similar, showing that tissue-specific eQTLs mostly do not reflect tissue-specific expression. Expression is defined as the median of log-Reads per Kilobase Mapped (log-RPKM) across individuals. Densities for genes having tissue-specific eQTLs (red) are estimated using average expression levels from *n* = 201–301 genes, depending on the tissue, and densities for remaining genes (black) are based on at least *n* = 15,768 average gene expression levels.

**Supplementary Figure 9.**
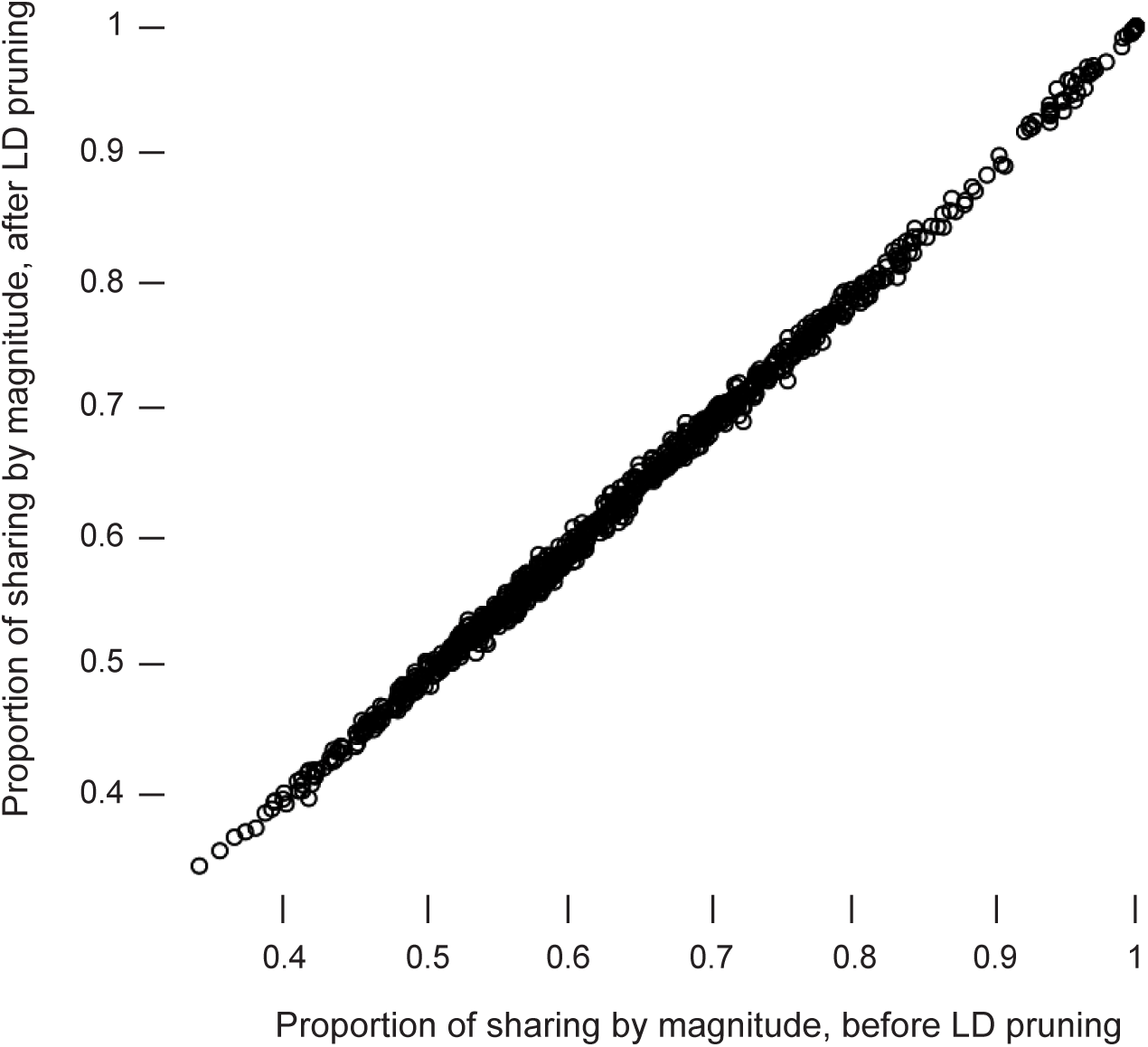
Comparison of pairwise sharing by magnitude for top eQTLs, without and with LD pruning. Each point corresponds to a pair of tissues. The horizontal axis gives results from the original `mash` analysis reported in the main paper; the vertical axis shows results from an “LD-pruned” analysis, where training data and top eQTLs were first pruned (using PLINK^12^) to avoid any pair of SNPs being in LD (*r*^2^ > 0.2) before `mash` was applied. The strong similarity of the results illustrates the robustness of `mash` to LD pruning.

**Supplementary Table 1.** Accuracy of effect size estimates for each method. Table shows relative root mean squared error (RRMSE) for all effects (RRMSE^All^), for the subsets of effects that are truly non-null (*β* ≠ 0; RRMSE^Non-null^) and truly null (*β* ≠ 0, RRMSE^Null^). RRMSE values less than 1 indicate improvements in accuracy over the original estimates. Values of RRMSE^Null^ < 1 indicate that shrinkage toward zero helped improve estimates of null effects. Values of RRMSE^Non-null^ < 1 indicate that pooling information across conditions can improve accuracy of estimates of non-null effects. Note that, in the “independent” simulations, most effects are null, so shrinkage of all methods improved overall performance compared to no shrinkage (RRMSE^All^ < 1) at the expense of lowering accuracy in the non-null effects (RRMSE^Non-null^ > 1). RRMSE^Non-null^, RRMSE^Null^ and RRMSE^All^ values were calculated from *n* = 400, 19,600 and 20,000 and observed effects, respectively, in 44 simulated tissues.

| method | Simulation framework | RRMSE<sup>All</sup> | RRMSE<sup>Non-null</sup> | RRMSE<sup>Null</sup> |
| --- | --- | --- | --- | --- |
| mash | shared, structured | 0.06 | 0.44 | 0.015 |
| mash-bmalite | shared, structured | 0.11 | 0.78 | 0.018 |
| ash | shared, structured | 0.21 | 1.34 | 0.076 |
| mash | shared, unstructured | 0.14 | 1.00 | 0.014 |
| mash-bmalite | shared, unstructured | 0.15 | 1.03 | 0.014 |
| ash | shared, unstructured | 0.21 | 1.37 | 0.078 |
| mash | independent | 0.28 | 1.82 | 0.112 |
| mash-bmalite | Independent | 0.28 | 1.82 | 0.118 |
| ash | Independent | 0.21 | 1.37 | 0.076 |

**Supplementary Table 2.** Overlap in associations identified from simulated data sets. Table summarizes the overlap in significant associations (*lfsr* < 0.05) identified among all methods that were compared. In both “shared effects” scenarios, `mash` captured the vast majority of the associations identified by the other methods. All association counts in the table are a subset of *n* = 20,000 × 44 = 880,000 simulated gene-SNP effects (most of which are zero).

| associations | —simulation framework— |  |  |
| --- | --- | --- | --- |
|  | shared, structured | shared, unstructured | independent |
| mash, not ash, not mash-bmalite | 3,889 | 622 | 32 |
| ash, not mash, not mash-bmalite | 0 | 0 | 740 |
| mash-bmalite, not mash, not ash | 37 | 9 | 79 |
| mash, ash, not mash-bmalite | 7 | 0 | 44 |
| mash, mash-bmalite, not ash | 5,777 | 336 | 70 |
| ash, mash-bmalite, not mash | 0 | 0 | 10 |
| all | 3,477 | 2 | 5,962 |

**Supplementary Table 3.** Tissue sample sizes. Right-hand column gives the sample size (*n*) for each tissue in the GTEx data set.

| **Tissue** | **sample size** |
| --- | --- |
| adipose visceral omentum | 227 |
| adrenal gland | 145 |
| artery aorta | 224 |
| artery coronary | 133 |
| artery tibial | 332 |
| brain anterior cingulate cortex BA24 | 84 |
| brain caudate basal ganglia | 117 |
| brain cerebellar hemisphere | 105 |
| brain cerebellum | 125 |
| brain cortex | 114 |
| brain frontal cortex BA9 | 108 |
| brain hippocampus | 94 |
| brain hypothalamus | 96 |
| brain nucleus accumbens basal ganglia | 113 |
| brain putamen basal ganglia | 97 |
| breast mammary tissue | 214 |
| cells EBV-transformed lymphocytes | 118 |
| cells transformed fibroblasts | 284 |
| colon sigmoid | 149 |
| colon transverse | 196 |
| esophagus gastroesophageal junction | 153 |
| esophagus mucosa | 286 |
| esophagus muscularis | 247 |
| heart atrial appendage | 194 |
| heart left ventricle | 218 |
| liver | 119 |
| lung | 320 |
| muscle skeletal | 430 |
| nerve tibial | 304 |
| ovary | 97 |
| pancreas | 171 |
| pituitary | 103 |
| prostate | 106 |
| skin not sun exposed suprapubic | 250 |
| skin sun exposed lower leg | 357 |
| small intestine terminal ileum | 88 |
| Spleen | 104 |
| Stomach | 193 |
| Testis | 172 |
| thyroid | 323 |
| uterus | 83 |
| vagina | 96 |
| whole blood | 393 |

**Supplementary Table 4.** Overlap in associations identified from GTEx data. Table summarizes the overlap in significant associations (*lfsr* < 0.05) identified among all methods compared. The `mash` method captures the vast majority of the associations identified by the other methods—only 248 associations identified by `ash` or `mash-bmalite` are not identified by `mash`—in addition to many other associations that are not identified by either `ash` or `mash-bmalite` (63,956). All association counts in the table are a subset of the *n* = 16,069 × 44 = 707,036 gene-SNP effects considered.

| associations | count |
| --- | --- |
| mash, not ash, not mash-bmalite | 63,956 |
| ash, not mash, not mash-bmalite | 2,383 |
| mash-bmalite, not mash, not ash | 11,789 |
| mash, ash, not mash-bmalite | 665 |
| mash, mash-bmalite, not ash | 176,572 |
| ash, mash-bmalite, not mash | 248 |
| all | 88,459 |

